# I. Phantom: a multiplex CRISPR-Cas9 gRNA toolkit for efficient generation of fully unpigmented F0 zebrafish

**DOI:** 10.64898/2026.10.26.761803

**Authors:** Axel Benchetrit, Arnim Jenett, Marjorie Juchaux, Jean-Pierre Levraud, Pierre Affaticati

## Abstract

Zebrafish are an increasingly popular model organism, valued for their natural transparency during embryonic stages. However, imaging at later developmental stages is affected by several types of pigments distributed over the body surface. These pigments partially or completely absorb or reflect light, and some also exhibit fluorescence, thereby interfering with imaging. Chemical approaches, such as phenylthiourea (PTU), are available to address this problem. However, PTU only inhibits melanin production and can affect development, making its use controversial. Many mutant zebrafish lines lacking single pigments have already been described, and combinations of melanin and iridophore deficient lines such as the *casper* and *crystal* fish greatly improved *in vivo* imaging at late stages. However, breeding multiple recessive mutations with a desired reporter transgene is laborious, and xanthophores remain in these lines. Here, we address these limitations by developing and optimising a multiplex CRISPR-Cas9 gRNA set that simultaneously targets the three major pigment types, while minimising mortality. This approach generates a new phenotype, termed *phantom*, characterised by the complete absence of visible pigmentation directly in F0 zebrafish larvae, while retaining the ability to survive to adulthood and reproduce.

## Introduction

The zebrafish (*Danio rerio*) has become a widely-used model organism in scientific research, mainly due to its accessibility, rapid development and natural transparency during its larval stages. However, as the organism matures, pigmentation increasingly obscures this transparency, making image-based phenotyping of specimens older than 1 day post-fertilisation (dpf) more difficult. The three distinct types of pigment cells found in the skin originate either directly from the neural crest or indirectly from hypodermal cell progenitors ^1^. In addition, pigment-producing cells are found in retinal pigment epithelium (RPE) that derive from the dorsal optic neuroepithelium ^2^.

**Melanophores**, which originate from the neural crest, are responsible for the irregular black spots along the dorsal region of larval zebrafish, extending from head to tail, as well as the characteristic black stripes seen in adulthood. Retinal pigment epithelium (RPE) cells also contain melanin, which accounts for the black pigmentation of the eyes. Melanin absorbs most wavelengths of visible light. Melanin deficiency can result from the inactivation of several genes, including *slc45a2*, which regulates melanosome acidification ^3^ and produces an albino phenotype when disrupted ^4,5^. A similar phenotype results from loss-of-function mutations in *tyr,* which encodes tyrosinase, an enzyme that converts tyrosine into DOPA-quinone during melanin synthesis in many vertebrates ^6,7^. Tyrosinase should not be confused with the tyrosine hydroxylases *th1* and *th2*, which produce L-DOPA for dopamine synthesis.

In contrast, the *nacre* phenotype results from *mitfa* knockout (KO), which specifically affects the population of cutaneous melanophores by preventing their differentiation from neural crest cells ^8–10^. Consequently, nacre mutants retain pigmented retinas but lack stripes along their flanks.

Chemical inhibition of melanin synthesis is also commonly achieved by addition of PTU^11^ (phenylthiourea, not to be confused with the antithyroid drug propylthiouracil). However this treatment has some developmental effects (Li, Z., Ptak, D., Zhang, L., Walls, E.K., Zhong, W. and Leung, Y.F., 2012. Phenylthiourea specifically reduces zebrafish eye size. *PloS one*, *7*(6), p.e40132.) and cannot be maintained for long periods of time, particularly when feeding of larvae becomes necessary.

**Iridophores** represent a second major population of pigment cells. They contain silvery guanine crystals that interfere with imaging by reflecting light and exhibit red autofluorescence. These silvery pigments are also present over the eyes from an early developmental stage. In larvae, iridophores are present in dorsal and ventral stripes and cover the swim bladder. In adults, they are found in the interstripes between melanophores and cover the abdomen and operculum. These cells help organise the cutaneous melanophores of the body into stripes, which explains why the *tra* and *roy* phenotypes, both caused by *mpv17* inactivation, are associated with a black-dot phenotype ^12,13^. Although spontaneous *mpv17* deficiency has been implicated in the sparsely pigmented roy phenotype, this mutation has several drawbacks. As *mpv17* encodes a widely expressed mitochondrial protein, its function is not restricted to iridophores but extends to other tissues. Consequently, spontaneous *mpv17* mutants are used as models of mitochondrial DNA depletion syndromes, while *mpv17* KO lines in zebrafish have been reported to exhibit abnormal muscle and liver development, as well as energy depletion phenotypes ^14,15^. Notably, *mpv17* KO zebrafish have a high mortality rate within the first four weeks of life due to developmental defects, whereas this is generally not observed with the spontaneous *mpv17/roy* phenotype ^12,13^ in zebrafish, mice ^14^ and humans ^18^. We hypothesise that this difference is related to the nature of the mutation. The spontaneous 19 bp deletion in *mpv17* is not located in an exon but in the second intron and may therefore cause an aberrant splicing instead of directly introducing a premature stop codon ^13^. However, mutations in *ltk*, whose expression is restricted to the neural crest and iridophores, have been shown to produce a very similar pigmentation phenotype ^19^. Interestingly, *ltk* loss-of-function also eliminates the silvery pigmentation over the retina. The silvery cells covering the eyes appear to originate from the neural-crest and are located on the surface of the RPE. Targeting *ltk*, therefore, offers an alternative approach to eliminate iridophore pigmentation, as also demonstrated in killifish^20^. To date, no effects on other tissues have been reported following ltk depletion.

**Xanthophores** produce a third, less-studied type of pigment, ranging from yellow to orange. They are distributed throughout the body and can be observed as early as two days post-fertilisation, mainly along the dorsal region of larvae. Unlike other pigment types, xanthophore pigmentation does not result from a single molecule but from a group of compounds known as pteridines, including sepiapterins and drosopterins, which accumulate within pterinosomes ^21^. These pteridines absorb light most strongly in the blue spectrum, the same wavelengths used to excite green fluorescent proteins such as eGFP (450-490 nm)^22^. This interferes with the acquisition of green fluorescence signals, at least in confocal microscopy. Furthermore, the spontaneous, uncharacterized *yobo* and *brie* mutations reduce yellow pigmentation in zebrafish, resulting in partial to near-complete loss of blue light absorbance, respectively ^22^.

Targeting transcription factors required for the differentiation of neural crest precursors into xanthophores, such as *pax7* or *csf1ra,* can disrupt yellow pigmentation. However, *pax7* is duplicated in the zebrafish genome as *pax7a* and *pax7b,* both of which must be knocked-out to prevent xanthophore differentiation ^23,24^. Furthermore, although pax7a/b-deficient zebrafish survive to adulthood, they exhibit muscle satellite cell loss and impaired regeneration, as do *pax7* KO mice ^25^. Loss of the receptor tyrosine kinase *csfr1a* produces the *panther* phenotype, in which the loss of xanthophores disrupts the melanophore stripe organisation ^26^. However, *csf1ra* knock-out is problematic because it increases locomotor activity without completely eliminating yellow pigmentation in adulthood ^27^. Moreover, *csf1ra* plays a role in macrophage differentiation and scar formation ^28–30^. Notably, *csfr1a* mutants lack microglia, the resident macrophages of the brain, during the larval stage ^31^. Knocking out this gene, therefore, has important side effects, particularly on the immune system.

However, in medaka (Oryzias latipes), loss of *slc2a11b* results in the absence of yellow pigmentation ^32^. Expression of this transporter appears to be restricted to pigment cells. Similarly, in *Xenopus tropicalis*, knock-out of its orthologue, *slc2a7*, has been shown to eliminate visible yellow pigmentation associated with xanthophores ^33^, providing a basis for developing more transparent animals ^34^.

Furthermore, *slc2a11b* zebrafish mutants generated by the Sanger Institute Zebrafish Mutant Project lack yellow pigmentation during larval development ^35^. We confirmed that the loss of *slc2a11b* is responsible for this phenotype by generating CRISPants in which reduced larval autofluorescence enabled the detection of faint green fluorescence associated with injected micelles^36^. Since no particular deleterious phenotype has yet been reported, the *slc2a11b* gene was targeted to produce our fully unpigmented phenotype.

Several zebrafish lines with pigment-loss phenotypes already exist. Single-gene mutants with pigmentation defects were first selected for the ornamental pet fish trade and have been known for many decades (Heffter 1996). More recently, these mutations have been combined to produce more transparent fish, particularly during adulthood. Combining the *nacre* (*mitfa*) and *roy* (*mpv17*) mutations produced the *casper* mutant^37^, which has become widely used for applications such as xenotransplantation in adults, despite its reduced fitness. However, the retina of *casper* fish remains black, prompting Antinucci *et al.* to develop the *crystal* phenotype by crossing *casper* double mutants with *slc45a2* (*albino*) mutants ^5^. Despite its improved transparency, this line has not become as widely adopted as *casper*, possibly because of the additional effort required to incross three separate recessive mutations within a transgenic background of interest. Furthermore, these fish retain xanthophores.

Consequently, generating transparent phenotypes through genetic engineering offers significant advantages over conventional breeding strategies. It can substantially reduce the number of generations required compared with conventional inbreeding, while also reducing the coefficient of consanguinity. Furthermore, the emergence of CRISPR-Cas9-based CRISPant techniques enables the simultaneous disruption of multiple loci, allowing complete phenotypes to be generated directly in F0 individuals. This eliminates the need to raise fish to adulthood before phenotypic analysis. We therefore aimed to generate zebrafish lacking all three pigment types directly in the F0 generation through multiplex gRNA-Cas9 injections, while maintaining acceptable mortality and malformation rates. Here, we report several strategies for generating *crystal*-like phenotypes, as well as a triple-pigment-less phenotype entirely devoid of melanin, iridophores and xanthophores, which we termed *phantom*.

## Results

### Choosing target genes by assessing the efficiency of individual and coupled knock-outs

The objective of this study was to identify combinations of gRNAs capable of completely abolishing pigment production in ≥50% of injected zebrafish larvae (Figure 1A), while maintaining low levels of developmental malformations associated with the pleiotropic effects of CRISPR-based injections.

**Figure 1:**
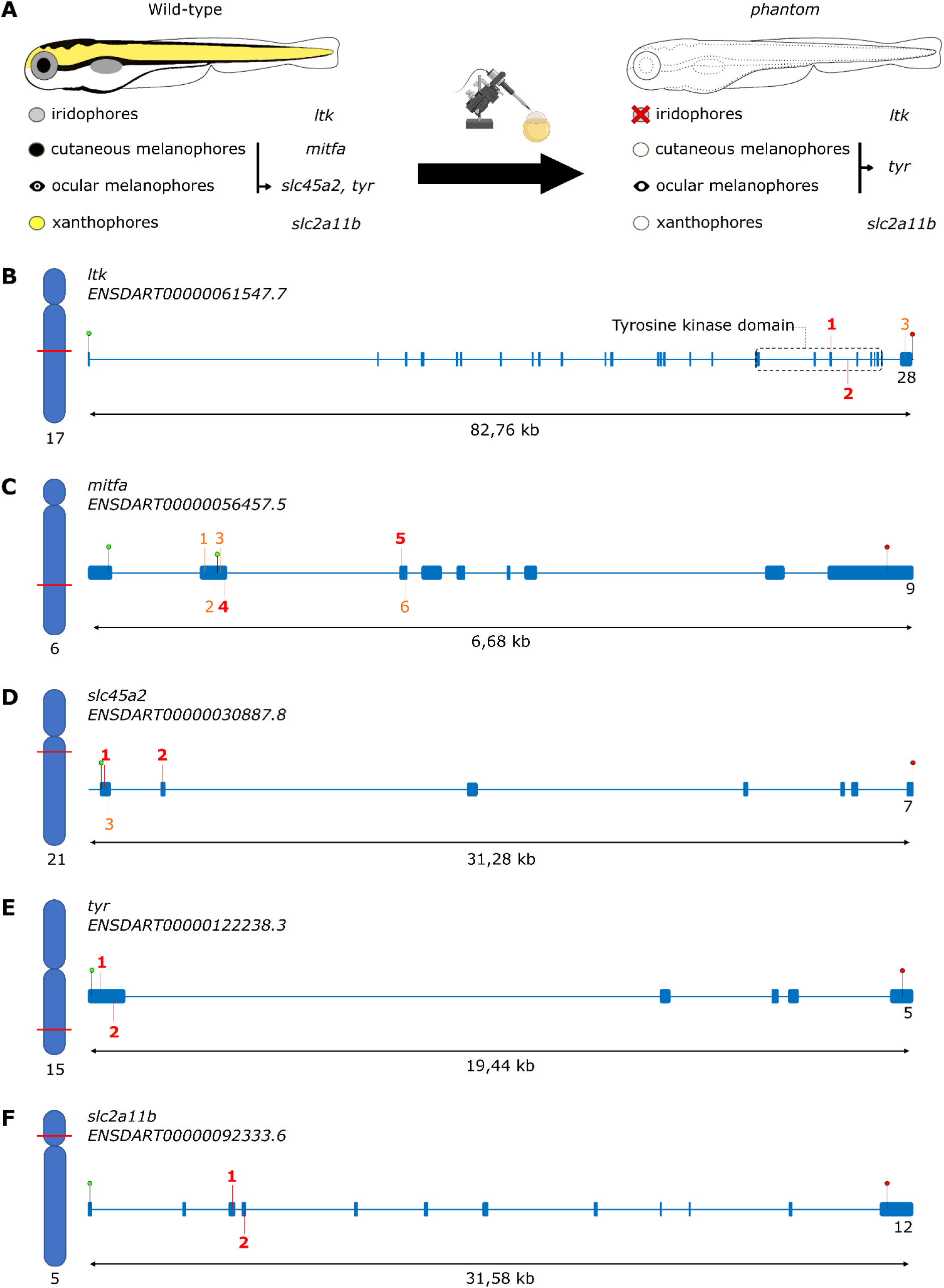
Generation of a phantom phenotype by multiplex CRISpants injections. 5 different genes (main annotated transcripts) are knocked-out using CRISPR-SpCas9: **(A)** pigments in zebrafish and their corresponding genes. While ltk disruption eliminates the iridophores, interference with the other genes only suppresses the pigment synthesis – either by disrupting the mechanism itself (tyr) or the importation of substrates needed for the pigment production (slc2a11b); **(B) ltk** main transcript is needed for the differentiation of neural crest cells into iridophores; **(C) mitfa** allows for the differentiation of neural crest cells into cutaneous melanophores. Two start codons are displayed to indicate the two potential main transcripts and the CRISPR strategy associated with this uncertainty. gRNAs for mitfa were chosen around the second start codon to ensure all main transcripts are hit; **(D) slc45a2** and **(E) tyr** are recruited (independently) for the production of melanin; The first is involved in transportation, and the second within the melanin synthesis. **(F) slc2a11b** serves in the transportation from the extracellular matrix to the cytosol for the production of sepiapterines in xanthophores. In each gene locus, displayed on its chromosome, green and red pins signal the start and stop codon of each gene. The locations of the tested gRNAs are labelled with their identification number. Two gRNAs per gene (shown in bold red) are selected to produce unpigmented larvae for each pigment. Among these 5 genes, 3 are selected to be knocked-out for the generation of the phantom phenotype: ltk, tyr, slc2a11b.

A total of 16 gRNAs targeting five genes were evaluated to identify efficient candidates for subsequent testing in pairs for each gene. These included three targeting *ltk*, six targeting *mitfa*, three gRNAs targeting *slc45a2* and two targeting *tyr* and *slc2a11b* each (Table 2). We designed three *slc45a2* gRNAs targeting the first and second exons to promote early premature stop codons in the single reported transcript of this gene (Figure 1D). As *mitfa* has two alternative transcripts encoding transcription factor domains, we selected gRNAs targeting exons 2–4 to ensure disruption of both transcripts (Figure 1C). For *tyr*, which has a single transcript, we targeted the first (and larger) exon (Figure 1E). For *ltk*, we designed gRNAs to disrupt the protein’s tyrosine kinase catalytic domain (Figure 1B). Finally, we selected *slc2a11b* gRNAs targeting exons 3 and 4, which are present in both protein-coding transcripts. These exons were chosen because of their proximity, increasing the likelihood of large deletions between the two cut sites (Figure 1F). All gRNAs were selected using CRISPOR ^38^ to minimise predicted off-target cleavage while maintaining on-target cutting efficiency (Table 2).

Among the ltk-targeting gRNAs, only ‘ltk 1’ induced substantial depigmentation, with ∼50% of larvae displaying a depigmented phenotype and <10% retaining wild-type pigmentation (Figure 2A). Although ‘ltk 2’, which targets an intronic region, and ‘ltk 3’, located in the last exon, produced high indel rates (Figure 1B), neither resulted in detectable depigmentation. These gRNAs were designed to promote large deletions between target sites when combined with ‘ltk 1’.

**Figure 2:**
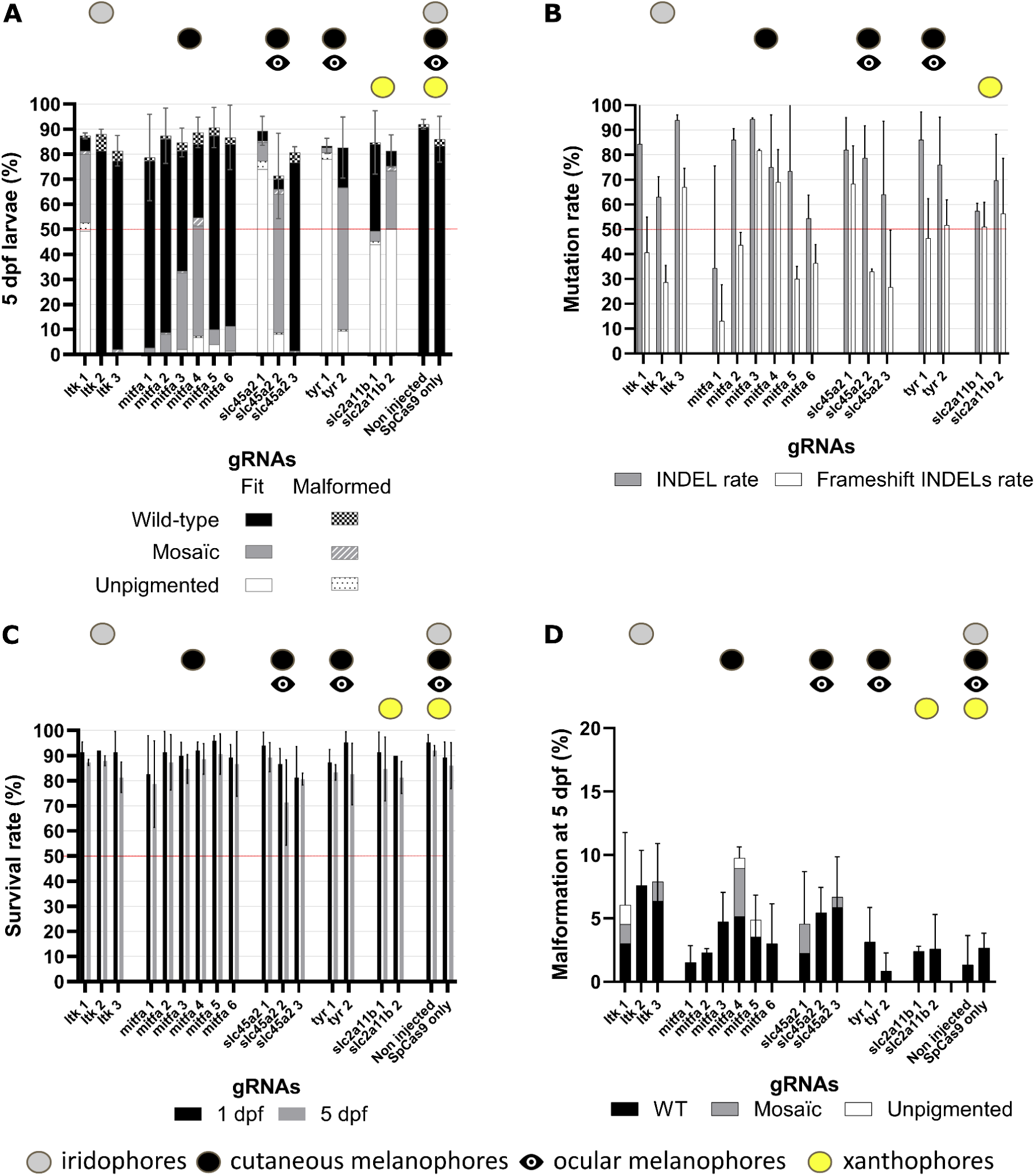
Characterisation of the effects of gRNA candidates on zebrafish larval development and pigment formation. **(A)** Histograms showing the percentage of 5 dpf larvae classified into six phenotype categories for each tested gRNA. Phenotypes were visually scored at 5 dpf and grouped into three pigmentation classes: no pigment loss (Wild-type, WT, black), partial pigment loss (Mosaic, grey) and complete pigment loss (Unpigmented, white). Each class was further subdivided into fit and malformed larvae (represented by checkered, striped and dotted patterns). **(B)** Total INDEL frequencies and frameshift-causing INDEL frequencies detected by Sanger sequencing of 1 dpf embryos prior to Synthego analysis, reported separately for each gRNA. **(C)** Histograms showing the percentage survival of zebrafish at both the embryonic and larval stages. The number of surviving individuals was recorded at 1 and 5 dpf. Negative controls (non-injected) and positive controls (SpCas9 only) were also assessed to ensure observed mortality was attributable to the gRNAs themselves. **(D)** Rates of malformations observed at 5 dpf for each tested gRNA. Every injection was subdivided into wild-type, mosaic and unpigmented phenotypes. For each gRNA, three independent batches of 50 one-cell stage zygotes (0 dpf) were injected and observed by a single observer. The red bar on the graphs highlights the 50% threshold we set as the minimum desired efficiency.

Injections targeting *mitfa* produced unexpected results. Only ‘mitfa 3’ and ‘mitfa 4’ generated mosaic depigmentation phenotypes in ∼30-50% of larvae. However, we observed almost no fully depigmented individuals (Figure 2A), despite high indel frequency and a substantial proportion of predicted frameshift mutations (Figure 2B). The lack of depigmentation following injection of ‘mitfa 1 and 2’ could be explained by the existence of an alternative *mitfa* transcript, ENSDART00000056457.5, with its start codon located just after the target sequences (Figure 1). This might suggest that this less-studied *mitfa* transcript contributes to melanocyte differentiation. However, this interpretation is complicated by the similarly weak effect of ‘mitfa 5 and 6’, which produced mosaic melanophore pigmentation in only 10% of larvae even though they target both transcripts.

The 50% threshold (red line in Figure 2, Figure 3, Figure 4) was chosen to ensure sufficient experimental throughput in F0 individuals. We selected several single-guide RNAs (sgRNAs; Table 1) for each target gene and injected them individually into one-cell-stage embryos (Figure 2). Cutting-efficiency and frameshift potential were assessed at each locus by deconvolution of mixed Sanger sequencing chromatograms using Synthego’s Inference of CRISPR edits (ICE) tool ^39^.

**Figure 3:**
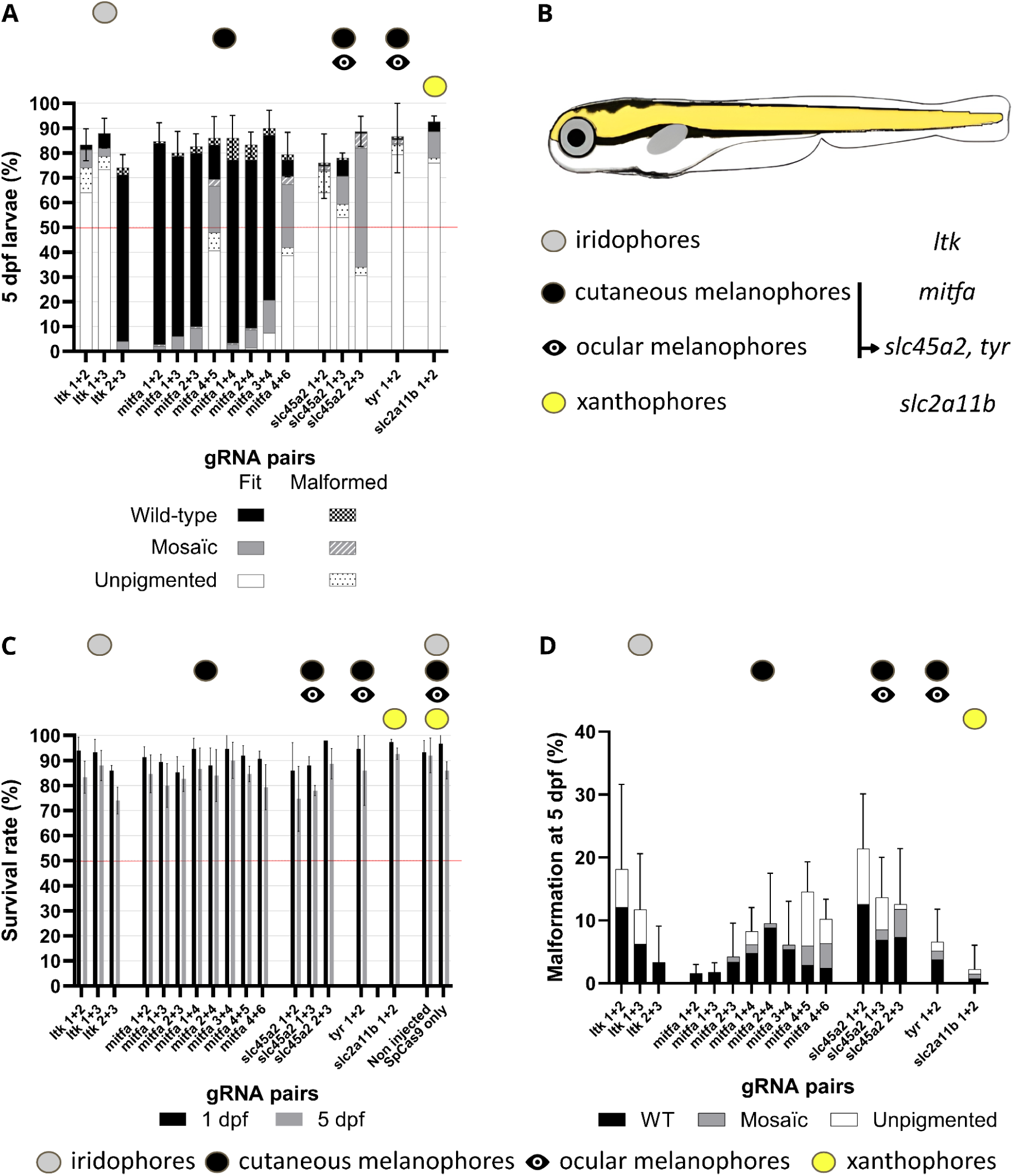
Characterisation of the effects of coupled gRNA candidates on zebrafish larvae pigment development. **(A)** Phenotypic ratios for each gRNA at 5 dpf, visually categorised into three groups: Wild-Type (WT, no pigment loss), Mosaic (incomplete pigment loss) and Unpigmented (total pigment loss). **(B)** Representation of a 5 dpf wild-type larva and pigment localisation related to targeted genes: iridophores (grey) are visible on the eye surface, around the lens and above the swim bladder; melanophores are subdivided into cutaneous (dorsal, ventral and midline) and ocular; yellow pigmentation is primarily dorsal above the midline. **(C)** Survival rates at 1 dpf (embryo) and 5 dpf (larval) stages; for each gRNA pair, 3 independent batches of 50 zygotes (0 dpf) were injected, with negative (non-injected) and positive (SpCas9 only) controls to confirm gRNA-specific mortality. **(D)** Malformation rates across gRNA injections and phenotypes; for each gRNA pair, three independent batches of 50 zygotes (0 dpf) were injected and scored by a single observer, with the red bar indicating the 50% minimum desired efficiency threshold.

**Figure 4:**
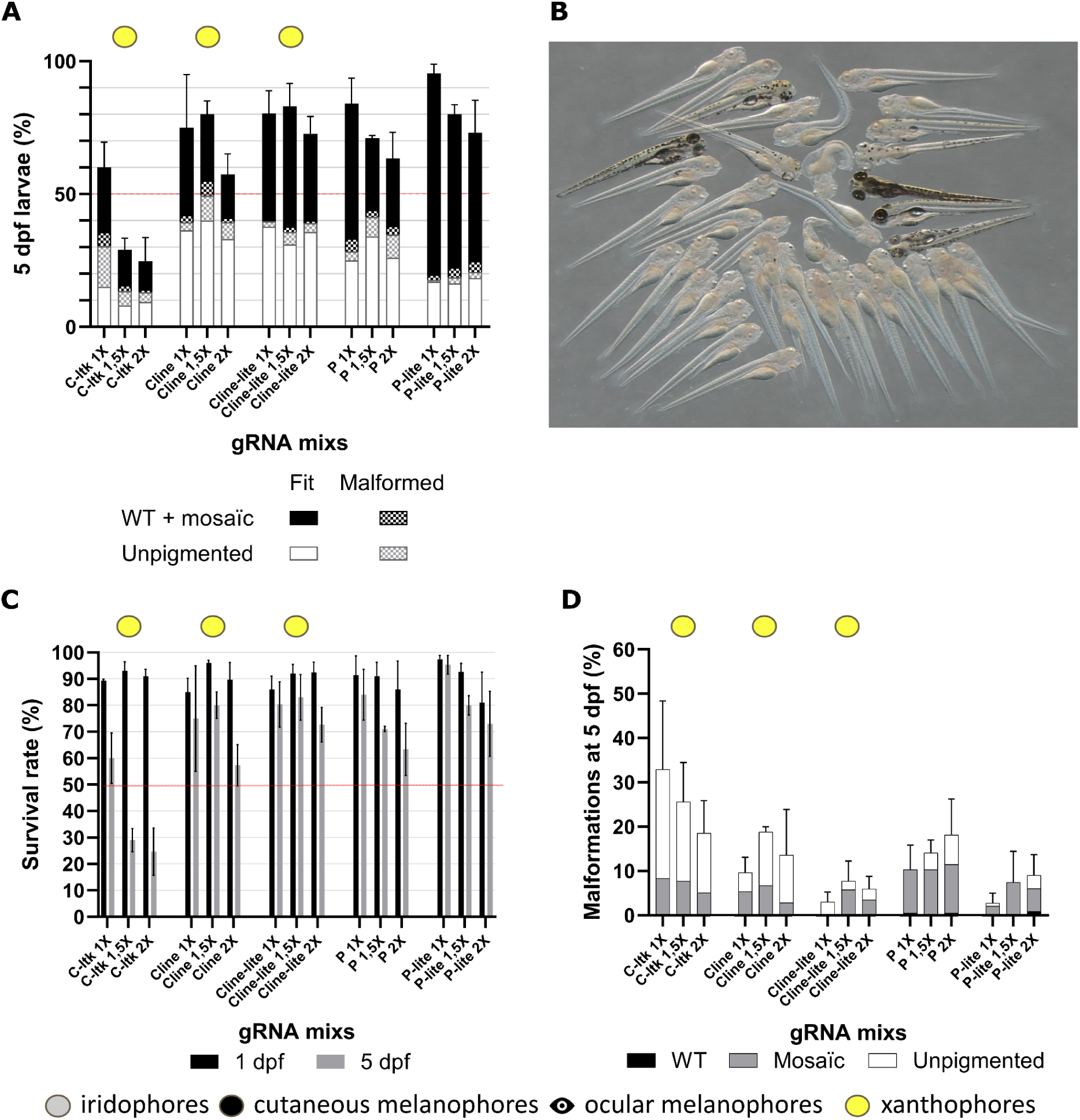
Characterisation of the effects of different gRNA mixes on zebrafish larval development to produce pigmentless phenotypes. **(A)** Phenotypic ratios for each gRNA at 5 dpf, visually categorised into two groups: pigmented (combining Wild-Type (WT) and Mosaic due to low WT frequency) and unpigmented (complete absence of black, yellow, or silvery pigment). **(B)** Degrees of mosaicism in a phantom 1.5× injection, where larvae were classified as unpigmented only if no pigment traces were visible. **(C)** Survival rates at 1 dpf (embryo) and 5 dpf (larval) stages; for each gRNA mix, 3 independent batches of 50 zygotes (0 dpf) were injected, with negative (non-injected) and positive (SpCas9 only) controls to confirm gRNA-specific mortality. **(D)** Malformation rates across gRNA injections and phenotypes; for each gRNA set, three independent batches of 100 zygotes (0 dpf) were injected and scored by a single observer, with the red bar indicating the 50% minimum desired efficiency threshold.

To validate the functional efficiency of each gRNA, as predicted from sequence analyses performed at 1 day post-fertilisation (dpf), we assessed larval pigmentation at 5 dpf (n = 50 larvae per condition; Figure 2. We classified larvae into three phenotypic categories according to the degree of pigment loss: wild-type, mosaic (partial disruption of the wild-type pigmentation pattern) and completely unpigmented (Figure 2D). Survival rates and morphological abnormalities were also recorded (Figure 2C).

Among the slc45a2-targeting gRNAs, ‘slc45a2 1’ produced the strongest phenotype, with *>70%* of larvae showing depigmentation (Figure 2A). In contrast, ‘slc45a2 2’ yielded ∼10% completely unpigmented larvae and <10% wild-type individuals, while most of the remaining larvae displayed mosaic pigmentation. ‘slc45a2 3’ produced no detectable depigmentation, consistent with sequence analyses showing very low frequencies of frameshift-inducing indels, despite an overall indel rate of 60–90%.

Embryos injected with ‘tyr 1’ displayed nearly complete depigmentation (∼100%), whereas ‘tyr 2’ predominantly generated mosaic larvae (∼70%) (Figure 2A).

Both *slc2a11b* gRNAs produced similar proportions of completely xanthophore-deficient larvae (∼50%). Notably, ‘slc2a11b 2’ yielded <5% wild-type individuals, compared with ∼40% following injection with ‘slc2a11b 1’.

Survival rates at 5 dpf *ranged from* 80–90% for all individual gRNA injections (Figure 2C), except for ‘slc45a2 2’, which showed a reduced survival rate of ∼70% with substantial variability. Overall, these survival rates were comparable to those observed in non-injected embryos and embryos injected with Cas9 alone. Among surviving larvae at 5 dpf, malformation rates ranged from 2-10%, regardless of pigmentation phenotype (Figure 2D). Injections of ‘mitfa 4’, as well as the *ltk* and *slc45a2* gRNAs, resulted in malformation rates >5%, whereas the remaining gRNAs produced frequencies similar to those observed in control embryos.

Interestingly, Sanger sequencing followed by ICE analysis showed only weak correlation with observed depigmentation efficiency for most gRNAs. This discrepancy was particularly evident for predicted frameshift rates, which were generally lower than expected from the phenotypic outcomes. An exception was observed for *slc45a2* gRNAs, for which sequencing results more closely matched depigmentation efficiency. In contrast, *mitfa* gRNAs showed medium-to-high in both cutting and predicted frameshift rates, despite producing minimal depigmentation.

As injections of individual gRNAs produced limited depigmentation for some target genes, particularly *mitfa*, we tested combinations of gRNAs to improve depigmentation by inducing large deletions between target sites and thereby disrupting gene function. To this end, we systematically paired the most efficient gRNA identified for each gene (‘ltk 1’, ‘mitfa 3’, ‘slc45a2 1’, ‘tyr 1’ and ‘slc2a11b 2’) with the remaining gRNAs targeting the same gene, including those that did not produce detectable depigmentation when injected individually. (Table 3).

For ltk, the combinations tested yielded relatively similar results (Figure 3). Pairing ‘ltk 1’ with another gRNA resulted in 60–70% complete depigmentation compared with ∼50% obtained with ‘ltk 1’ alone. The combination of ‘ltk 1 and ltk 2’ produced fewer iridophore-deficient larvae, but also fewer wild-type individuals. As expected, the ‘ltk 2 and 3’ combination resulted in almost no depigmentation, highlighting the importance of inducing targeted DNA cleavage early within the tyrosine kinase catalytic domain (Figure 3A). Because ‘ltk 2’ is located closer to ‘ltk 1’ than ‘ltk 3’, we selected the ‘ltk 1 and 2’ pair to generate the artificial *crysta-ltk* and *phantom* phenotypes. Interestingly, the swim bladder is not visible in larvae displaying the *ltk* CRISPant phenotype (Figure 5G), most likely due to the absence of light reflection, although normal swimming behaviour was observed. This observation warrants further investigation.

**Figure 5:**
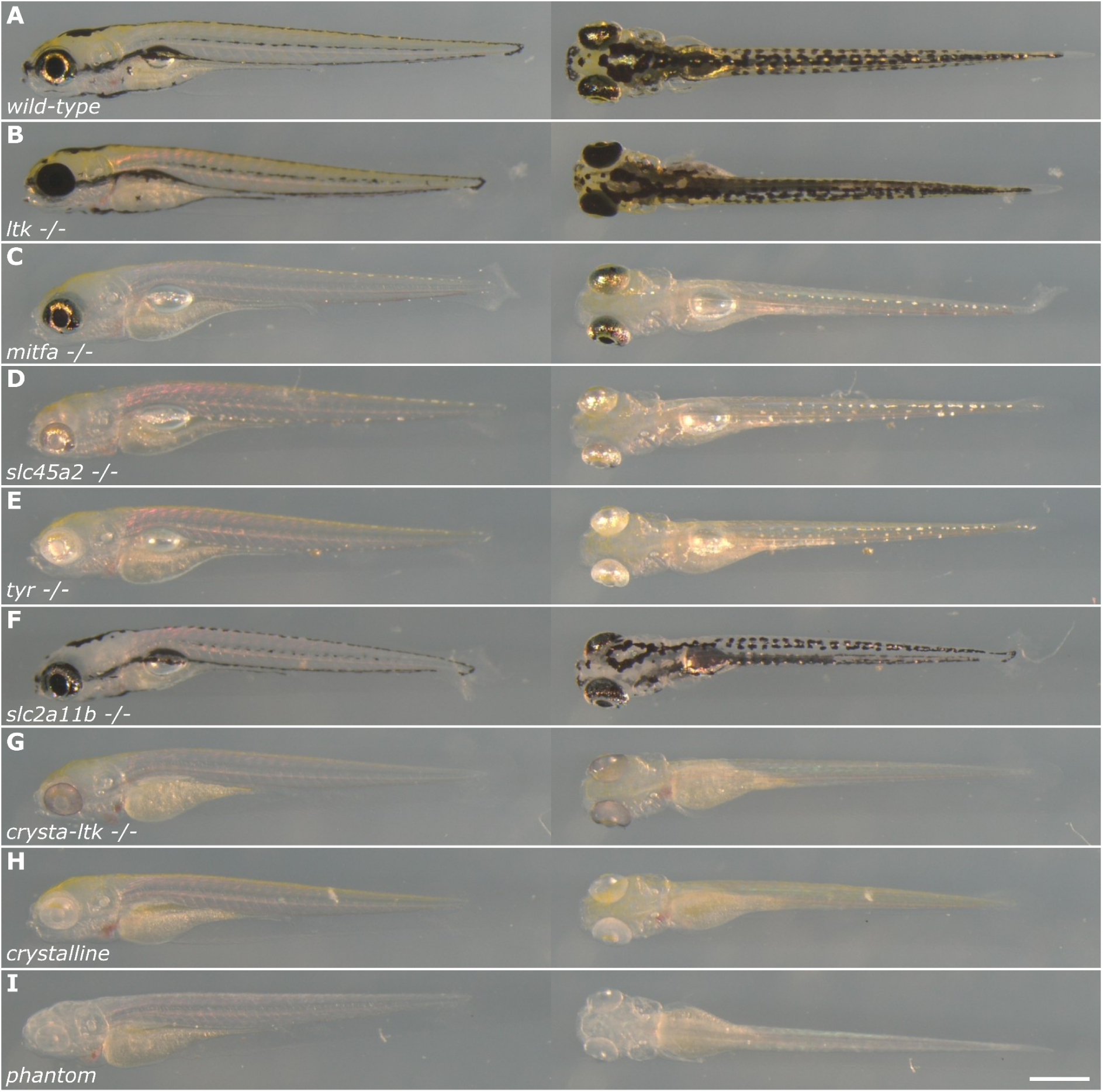
Mosaic of all depigmentation phenotypes at 5 dpf. Representative phenotypes are shown in lateral (left panel) and dorsal (right panel) views under a stereomicroscope with reduced-upright-angled bright field illumination to enhance visualisation of iridophores. (**A**) The wild-type larva has all three pigments : melanophores in the dorsal and ventral part of the body and in the lateral line and eyes, iridophores on the outer layer of the eye and along the body and xanthophores above the lateral line. Single depigmentation phenotypes were generated by the most efficient coupled gRNA injections for each gene (**B–F**: ltk, mitfa, slc45a2, tyr, slc2a11b) as well as for the combined crysta-ltk, crystalline and phantom phenotypes (**G–I**). Scale bar: 0.5 mm.

Surprisingly, although ‘mitfa 4’ produced ∼40% complete depigmentation when paired with either ‘mitfa 5’ or ‘mitfa 6’, ‘mitfa 3’ was less efficient when combined with ‘mitfa 1’ or ‘mitfa 2’ than when injected alone. Similarly, ‘mitfa 4’ showed reduced efficiency in these combinations compared with its individual performance. The remaining combinations produced almost no depigmentation.

As expected from the individual gRNAs, the ‘slc45a2 1 and 2’ combination was the most efficient among the *slc45a2* pairs (95%), followed by ‘slc45a2 1 and 3’ (70%) and ‘slc45a2 2 and 3’ (40%) (Figure 3A). In completely depigmented larvae, the loss of cutaneous melanophore pigmentation appeared comparable to that observed following *mitfa* knockout, but *slc45a2* disruption also resulted in eye depigmentation.

Similarly, the *tyr* pair produced 95% black depigmentation. Interestingly, *tyr* knock-out appeared to depigment the eyes more efficiently than *slc45a2* knock-out (Figure 3). This difference was also observed in adults. Nevertheless, adult F0 knock-out phenotypes were similar for both *tyr* and *slc45a2*, showing faded pigment stripes (Figure 5).

The slc2a11b pair resulted in the loss of yellow pigmentation in 80% of larvae (Figure 3A). This improvement likely reflects the moderate efficiencies of the two individual gRNAs.

With the exception of the ‘slc45a2 1 and 2’ and ‘ltk 2 and 3’ combinations, which showed ***∼***30% mortality at 5 dpf, all gRNA pairs produced survival rates >80% (Figure 3). However, the combinations ‘ltk 1 and 2’, ‘slc45a2 1 and 2’, ‘mitfa 4 and 5’ and ‘mitfa 4 and 6’, which were also the most efficient pairs for their respective genes, resulted in higher malformation rates of ∼15-20% *at 5 dpf* (Figure 3).

Based on these results, we aimed to generate CRISPant larvae that phenocopy the original *crystal* zebrafish model, phenotypically characterised by the absence of black and silver pigmentation due to deficiencies in *mitfa*, *slc45a2* and *mpv17.* To achieve this, we engineered a modified genetic combination termed *crysta-ltk*, in which *mpv17* was substituted with *ltk* (‘ltk 1 and 2’, ‘slc45a2 1 and 2’, ‘mitfa 4 *and 5’*). To improve survival rates and targeting efficiency, we then developed an alternative gRNA set, termed *crystalline, containing fewer* guides (‘ltk 1 and 2’, ‘tyr 1 and 2’). A further simplified configuration, *crystalline*-lite (‘ltk 1’, ‘tyr 1’), was designed to reproduce the *crystalline* phenotype while minimising off-target effects and improving viability. To generate the *phantom* phenotype, which lacks all three pigment types, we expanded the *crystalline* set by adding the *slc2a11b* pair (‘slc2a11b 1 and 2’). Finally, we established a corresponding *phantom*-lite configuration (‘ltk 1’, ‘tyr 1’, ‘slc2a11b 2’) to assess whether reducing the number of guides could improve the balance between depigmentation efficiency and survival (Table 4).

### Generation and assessment of the depigmentation capacity of the artificial *crystal* and pioneer *phantom* phenotypes

Three concentrations of *crysta-ltk* (C-ltk), *crystalline* (Cline) and *phantom* (P) gRNA sets were injected, ranging from 1X to 2X, in order to compensate for the expected efficiency loss caused by dilution between multiplexed gRNAs.

The *crysta-ltk* set showed a marked decrease in survival as gRNA concentration increased. Survival dropped from 60% to 25%, while malformation rates decreased from 30% to <20%. 30-50% of surviving larvae displayed a complete loss of pigmentation (Figure 4).

In comparison, the *crystalline* set yielded a better survival rate; ∼75-80% decreasing to 60% only at the 2X concentration, similar to *crysta-ltk* at 1X. Malformation rates remained substantially lower (10-18%), while most larvae still showed complete pigment loss. This improved outcome could be explained by both the lower toxicity of the injected gRNAs and the reduced number of gRNAs in the *crystalline* set (four vs six in *crysta-ltk*). As expected, increasing gRNA concentrations increased the proportion of fully depigmented larvae, although the total number of such larvae did not increase due to reduced survival. *Crysta-ltk* and *crystalline* phenotypes were largely indistinguishable except for eye depigmentation (Figure 5). Consistent with observations from injections of the ‘slc45a2 1 and 2’ pair compared with the ‘tyr 1 and 2’ pair, the eye appeared more pigmented in *crysta-ltk* larvae than in *crystalline* larvae (Figure 13, supp. 4). In both sets, *ltk* knock-out efficiency remained the limiting factor, silvery cells accounting for 50%-80% of the residual mosaic pigmentation. Notably, the stripes were absent in the artificial *crystal* phenotypes generated with *crysta-ltk* and *crystalline* sets (without melanocyte depletion), likely due to the absence of iridophores required for proper melanophore stripe organisation ^40^. Indeed, the classical *crysta-ltk* phenotype generated by *mitfa*, *slc45a2* and *ltk* knock-outs is nearly indistinguishable from the *crystalline* phenotype produced by *tyr* and *ltk* knock-outs at both larval and adult stages. The only difference lies in the eye, where *tyr* knock-out is better at removing the pigment than *slc45a2*.

As observed in *ltk*-CRISPants alone, the swim bladder was not visible in *crysta-ltk*, *crystalline* and *phantom* larvae, despite their normal swimming behaviour. However, the swim bladder was present in adults (Supplementary Fig. 7). This lack of a swimming bladder at 5 dpf could be a delay in its inflation or development for an unknown reason, since none of the targeted genes is involved in these processes. Several off-targets are located in the coding sequence of genes responsible for swimming bladder development (*ltk* gRNA 2: *diexf*, *slc45a2* gRNA 3: *asna1*) or inflation (*slc45a2* gRNA 1: *ca5a*, *mitfa* gRNA 6: *per2*). However, these genes are a concern mainly in *crysta-ltk* gRNA mixes and have 4 mismatches compared to the desired target. Consequently, these off-targets are very unlikely to be cut frequently and, therefore, cannot explain this “missing swimming bladder” phenotype amongst all *crystal* and *phantom* phenotypes.

The addition of the ‘slc2a11b 1 and 2’ pair to the *crystalline* set to generate the *phantom* set did not reduce survival to the same extent when concentrations increased. Survival decreased only from 80% at 1X to 70% at 2X, compared with 60% for the *crystalline* set. This effect may result from dilution of the ‘ltk 1 and 2’ gRNAs, thereby reducing their deleterious effects previously observed when injected alone (Figure 3) or combined only with ‘*tyr* 1 and 2’. Maximum efficiency for the *phantom* set was obtained at 1.5X rather than 2X, reaching ***∼***50%, with malformation rates comparable to those observed in the *crystalline* set (10%-15%). However, mosaic larvae were more frequent than fully depigmented larvae (Figure 4C). Based on these results, both *crystalline* and *phantom* sets reached the efficiency threshold defined at the beginning of the study.

To improve editing efficiency without further compromising survival, we tested reduced gRNA sets for both *crystalline* and *phantom*, termed *crystalline*-lite (Cline-lite) and *phantom*-lite (P-lite). These sets contained only the most efficient gRNA per gene (‘*ltk* 1’, ‘*tyr* 1’ and ‘*slc2a11b* 2’). In the *crystalline*-lite set, the proportion of fully depigmented larvae decreased compared with *crystalline*; however, the improved survival rate—particularly at 2X—compensated for this reduction, resulting in a similar total number of depigmented larvae as obtained with *crystalline* 2X. Malformation rates were also markedly reduced across all concentrations (<10%), likely reflecting fewer off-target effects and fewer cutting events.

In contrast, the *phantom*-lite set did not show a comparable improvement. Although malformation rates remained below 10% and survival rates were the highest observed among all tested sets (95% at 1X, 80% at 1.5X and 70% at 2X), editing efficiency decreased substantially. Only 15%-25% of larvae exhibited complete depigmentation across concentrations, suggesting excessive dilution of gRNAs and consequently fewer cutting events. Interestingly, at 1.5X and 2X concentrations, xanthophore depigmentation was the most affected phenotype, whereas *ltk* knock-out remained the limiting factor in all other *phantom* and *phantom*-lite conditions, regardless of concentration. Mosaic larvae from the P-lite 1X condition displayed a broad range of pigmentation patterns, reflecting the low editing efficiency at this concentration and highlighting the importance of the second ‘*tyr* 2’ gRNA in melanophore depigmentation despite its moderate knock-out rate (Figure 10 supp 1).

Overall, these results highlight the importance of increasing the number of cutting events rather than simply increasing the concentration of a smaller subset of gRNAs to maximise multiplex knock-out efficiency. As expected, this strategy reduces survival rates; however, the absolute number of usable (unpigmented) larvae is ultimately more critical than their proportion within the injected population. Consequently, *crystalline* 1.5X provides a similar yield of usable larvae as *crystalline*-lite at 1X, 1.5X, or 2X, albeit with slightly higher malformation rates. By contrast, *phantom* 1.5X appears to be the most suitable condition when efficient xanthophore depigmentation is desired.

### Enhancement of signal-to-noise ratio using *phantom* larvae

Reporter transgenes generated by inserting a fluorescent protein at the endogenous locus (“knock-ins”) are more reliable, but often substantially dimmer than those selected by Tol2-mediated random transgenesis, including a promoter sequence, for which the brightest lines that get chosen are probably those with insertion at a locus with minimal negative regulation. One strategy to overcome this limitation is the use of brighter, next-generation fluorescent proteins such as mScarlet3 or mNeonGreen. However, tagging genes with low endogenous expression levels remains challenging even with these improved reporters because their expression remains constrained by native regulatory elements. Although red fluorophores are generally preferred for detecting weak signals due to their improved signal-to-noise ratio compared with green fluorophores, we sought to investigate the contribution of xanthophores to background fluorescence when screening weak signals in both spectral channels. Xanthophores are known to exhibit autofluorescence when excited by blue light, potentially interfering with fluorescence detection ^22^.

To address this question, we crossed the lines *Tg(mpeg1:eGFP-CAAX)* and *Tg(mfap4.1:mCherryF)* to generate double-transgenic embryos. The lines display similar macrophage expression patterns with a relatively dim fluorescence in the green and red channels, respectively. To evaluate the effect of xanthophores on both eGFP and mCherry signals, *slc2a11b* gRNAs 1 and 2 were injected into one-cell stage *Tg(mpeg1:eGFP-CAAX);Tg(mfap4.1:mCherryF)* embryos. Fluorescence was assessed at 5 dpf in both channels.

In *slc2a11b* knock-out larvae, macrophages located in the dorsal region became readily detectable. In contrast, in wild-type larvae, these cells could not be distinguished from endogenous autofluorescence. The improvement in signal-to-noise ratio was particularly pronounced in the green channel: in wild-type larvae, autofluorescence was particularly visible above the midline due to xanthophore autofluorescence, whereas this signal was absent in *slc2a11b* knock-out larvae. In contrast, background fluorescence is barely detectable in the red channel of wild-type larvae, although it appeared slightly reduced compared with the knock-out condition (Figure 6).

**Figure 6:**
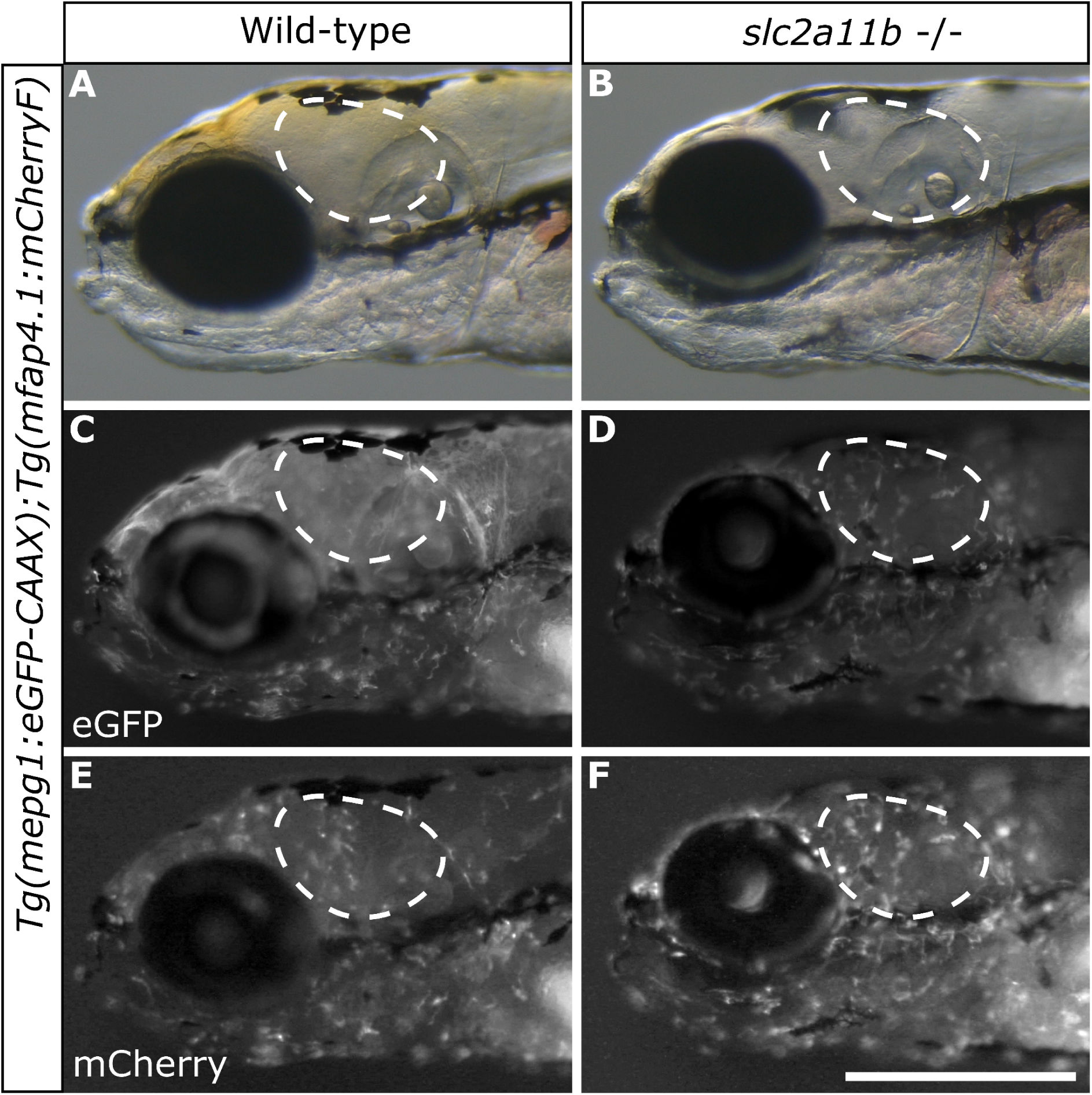
Qualitative comparison between Tg(mpeg1:eGFP-CAAX);Tg(mfap4.1:mCherryF) zebrafish injected with slc2a11b-targeting gRNAs or non-injected (heads). The Tg(mpeg1) line was crossed with the Tg(mfap4.1) line to generate double transgenic embryos, which were either injected at the 1-cell stage with slc2a11b **(B, D, F)** or left non-injected as wild-type controls **(A, C, E)**. At 5 dpf, larvae displaying expected wild-type and slc2a11b phenotypes were sorted and imaged under a fluorescence stereo microscope to assess the impact of xanthophore autofluorescence on macrophage pattern visibility in both the green **(eGFP-CAAX, C and D)** and red **(mCherryF, E and F)** channels. A white dashed area highlights an upper body region where xanthophore autofluorescence most strongly interferes with macrophage fluorescence detection. Brightness and contrast were uniformly optimised for both green and red channel images. Scale bar: 0.5 mm.

Interestingly, xanthophore-derived background fluorescence had an even stronger impact in the tail region, where macrophages were largely undetectable except in the most ventral part of the tail. Removal of xanthophore pigmentation improved the detection of mCherry fluorescence as well, although the effect was less pronounced than for the eGFP signal. Consistent with these observations, injection of the *phantom* gRNA set in the same genetic background enabled more reliable detection of both eGFP and mCherryF signals compared with the *crystalline* injections (Figure 7).

**Figure 7:**
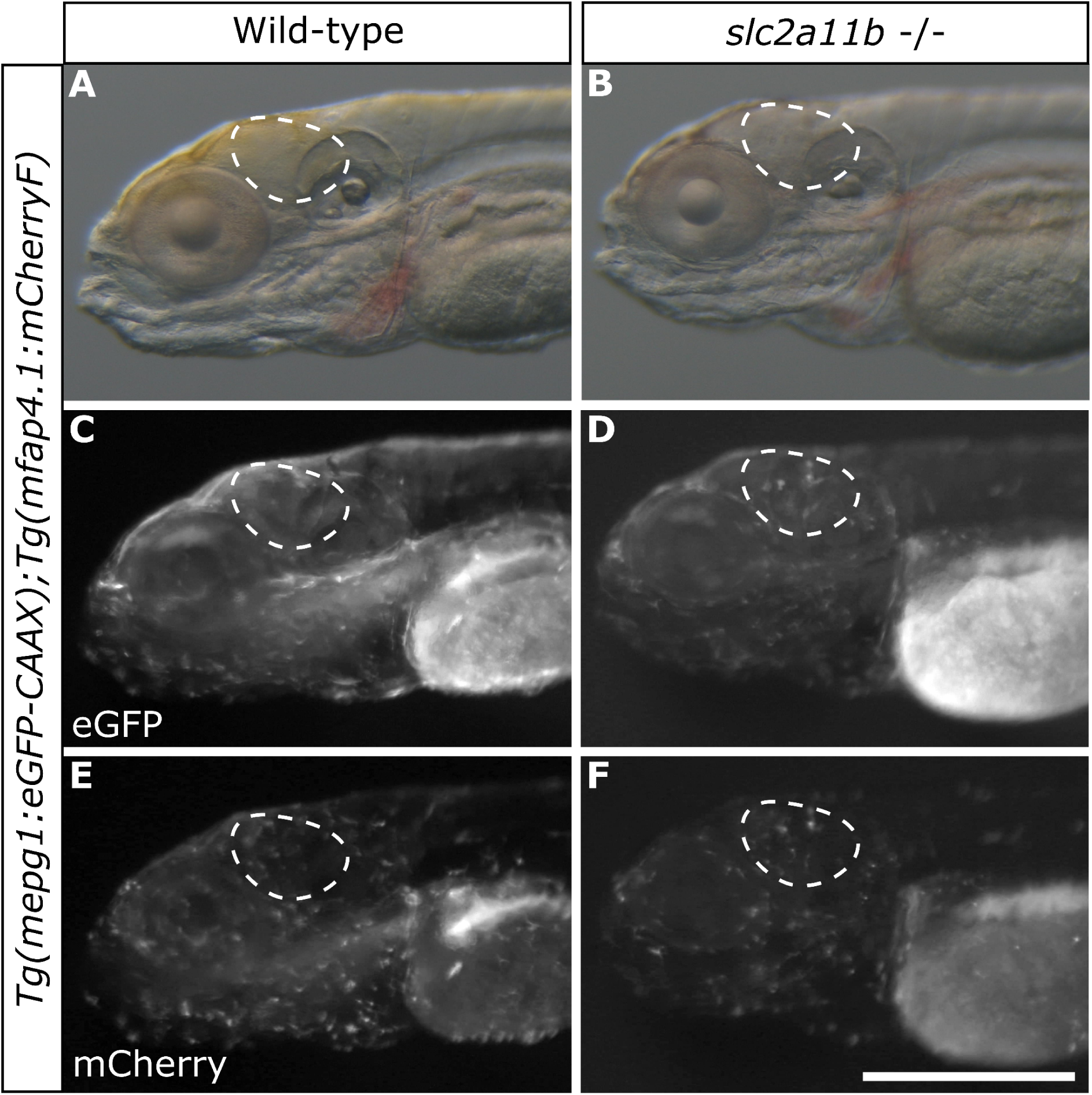
Qualitative comparison between Tg(mpeg1:eGFP-CAAX);Tg(mfap4.1:mCherryF) zebrafish injected with crystalline or phantom gRNA sets (heads). The Tg(mpeg1) line was crossed with the Tg(mfap4.1) line to generate double transgenic embryos, which were either injected at the 1-cell stage with crystalline **(A, C, E)** or the phantom gRNA set **(B, D, F)**. At 5 dpf, larvae displaying expected crystalline or phantom phenotypes were sorted and imaged under a fluorescence stereo microscope to assess the impact of xanthophore autofluorescence on macrophage pattern visibility in both the green **(eGFP-CAAX, C and D)** and red **(mCherryF, E and F)** channels. A white dashed area highlights an upper body region where xanthophore autofluorescence most strongly interferes with macrophage fluorescence detection. Brightness and contrast were uniformly optimised for both green and red channel images. Scale bar: 0.5 mm.

### Genetic depigmentation allows for 3D confocal imaging without immunostaining

We then sought to confirm the enhanced signal-to-noise ratio observed in live 5 dpf larvae under a stereomicroscope using confocal imaging after fixation and clearing. Typically, larvae are fixed in 4% PFA, followed by depigmentation via H₂O₂ treatment prior to incubation in the clearing medium. This H₂O₂ step removes endogenous fluorescence, thereby necessitating relabeling of the treated larvae by immunohistochemistry. With the *crystalline* and *phantom* gRNA sets, H_2_O_2_ treatment is not needed anymore, which allows for the acquisition of endogenous fluorescence. Moreover, the samples are less prone to physical damage during prolonged labelling procedures, thereby better preserving their structural integrity during handling and mounting.

We assessed the light penetration of genetically depigmented *crystalline* and *phantom* larvae. Therefore, we injected AB embryos with *crystalline* and *phantom* gRNA mix before staining them with YOYO-1 and DiIC18 to label the nuclei and the lipids (cell membranes). Together, these dyes give a general overview of the larval anatomy critical to ensure the light penetration throughout the entire specimen in both depigmented phenotypes. We compared them to H_2_O_2_-treated and untreated AB prior to tissue clearing in a dorsal-oriented acquisition before projecting them in lateral (Figure 8). As expected, AB larvae showed a strong depth-dependent gradient of signal with complete loss of signal under the dorsal cutaneous melanophores (Figure 8A and C). H_2_O_2_-treated larvae showed the best penetration, allowing for the imaging of the entire stack, still with a signal gradient. Both *crystalline* and *phantom* phenotypes showed the same penetration levels comparable to H_2_O_2_-treated larvae (Figure 8F and G) without deformation.

**Figure 8:**
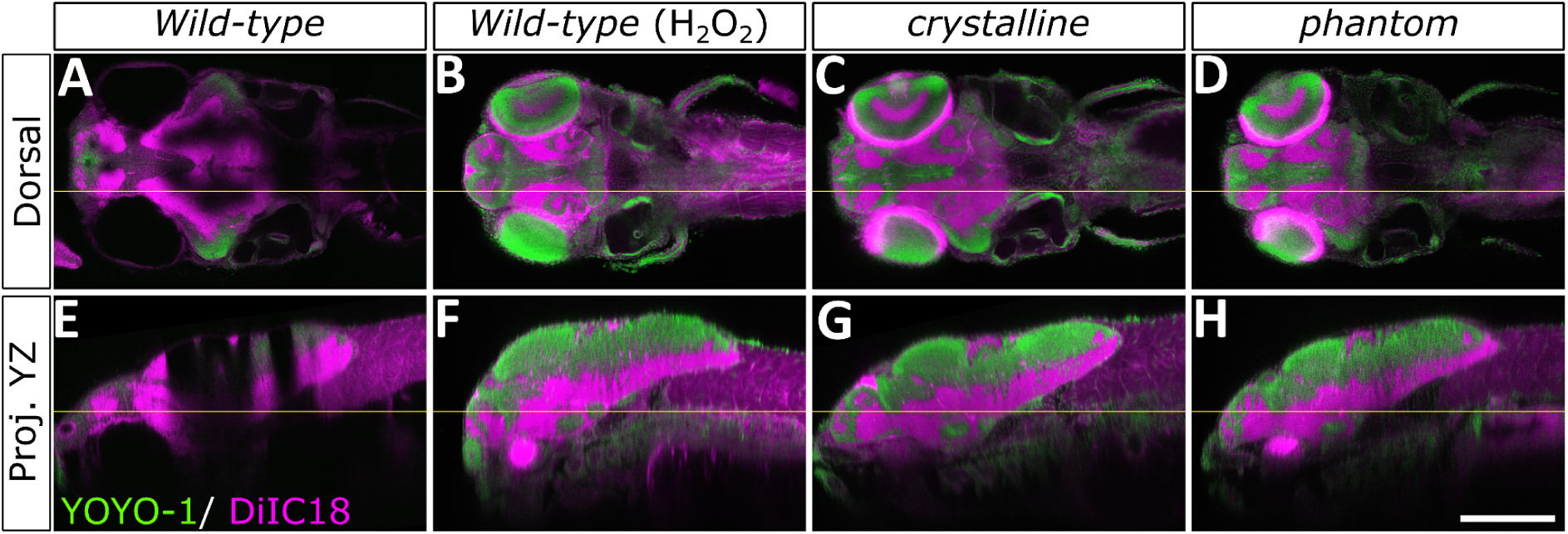
Comparison of light penetration between non-depigmented and depigmented larvae. 5 dpf wild-type larvae were fixed in 4% PFA before tissue clearing, then labelled with YOYO-1 (green) for nuclei and DiI-C18 dyes (magenta) for lipids (cell membranes and myelin sheaths). Imaging was performed under a Nikon A1 confocal microscope in dorsal orientation **(A–D)**. Acquisitions were later projected onto the lateral view **(E–H)** to assess light penetration and signal loss. Wild-type AB larvae without chemical or genetic depigmentation **(A, E)** exhibited drastically reduced light penetration compared to H₂O₂-treated larvae **(B, F)**, along with complete loss of fluorescent signals beneath dorsal melanophores. Crystalline **(C, G)** and phantom **(D, H)** phenotypes demonstrated light penetration levels comparable to H₂O₂-treated larvae **(B, F)**, without the fragility associated with chemical treatment, thus preserving body structure integrity after handling. Scale bar: 0.5 mm.

We then initially injected the two transgenic lines *Tg(mpeg1:eGFP-CAAX)* and *Tg(mfap4.1:mCherryF)* with *crystalline* and *phantom* gRNA mix, but the weak endogenous fluorescence was not observable anymore after the fixation step. This demonstrates one limit of endogenous fluorescence imaging. Instead, we used two other transgenic lines, *Et(slc18a2:eGFP)* and *Tg(kdrl:mCherry)*, with *crystalline* and *phantom* gRNA sets and compared them with their uninjected siblings (Figure 9). In both the green (Figure 9B and C) and red (Figure 9E and F) fluorescence channels, F0 depigmentation enabled the detection of numerous cells and structures, particularly within and beneath the eye tissues, as well as along the midline, where signal detection was otherwise hindered by wild-type pigmentation (Figure 9A and D).

**Figure 9:**
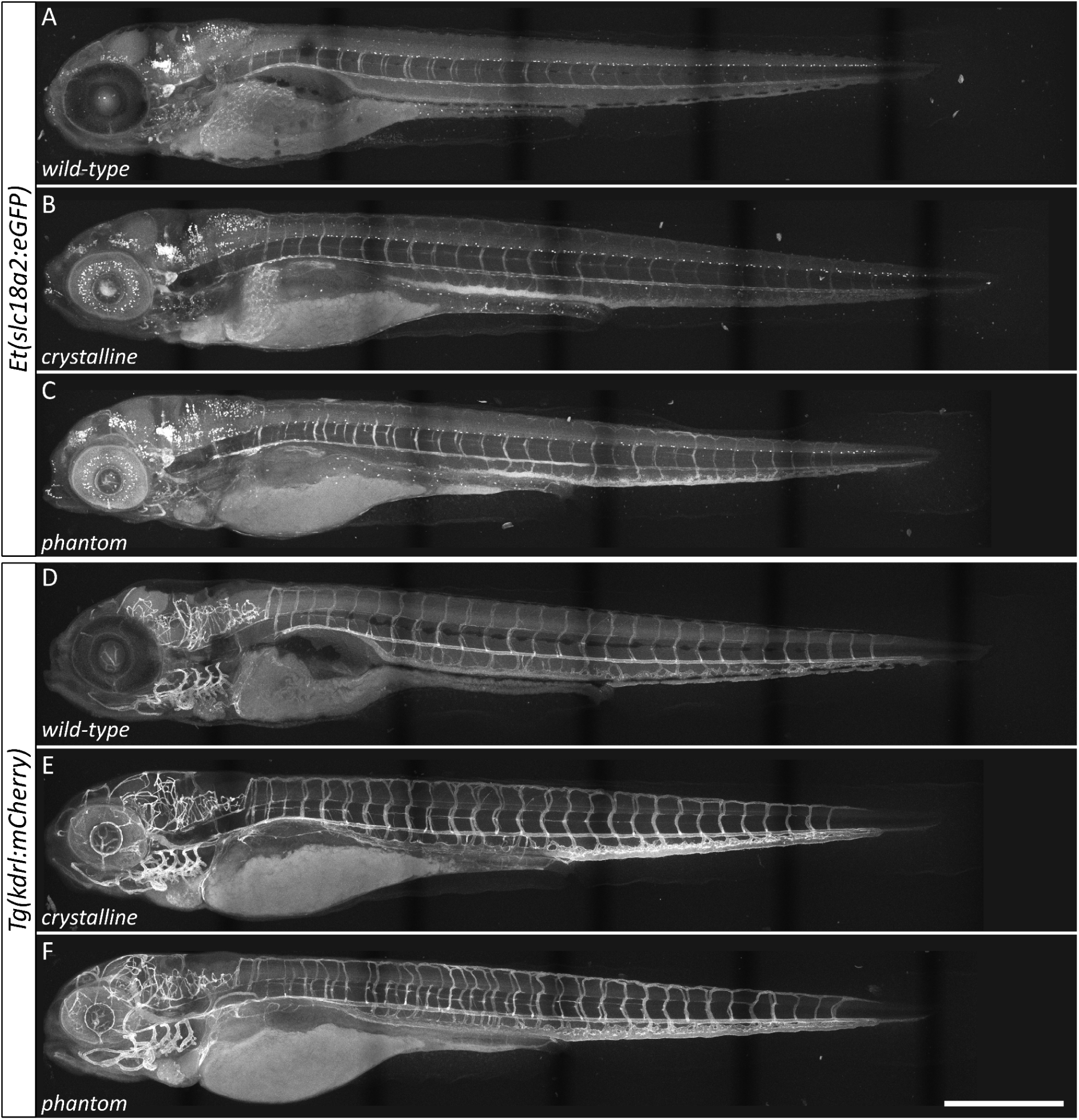
Comparison of confocal acquisition of transgenic lines depending on the depigmentation. 5 dpf Et(slc18a2:eGFP) **(A–C)** and Tg(kdrl:mCherry) **(D–F)** larvae were either non-injected **(A, E)** or injected with crystalline 1.5X **(B, E)** or phantom 1.5X **(C, F)**. They were fixed in 4% PFA before clearing and lateral confocal imaging. All larvae were imaged in the same session using identical parameters for green fluorescence **(A–C)** and red fluorescence **(D–F)**. While wild-type larvae **(A, E)** show no signal below melanophore-rich areas (eye and midline), both crystalline **(B, E)** and phantom **(C, F)** larvae exhibit identical signal patterns and intensity levels in both green and red channels. Scale bar: 0.5 mm.

However, we did not observe any difference between *crystalline* and *phantom*, especially in the upper body, where the xanthophores lie. When looking at the larvae in brightfield, we discovered that the yellow pigmentation, along with the reflective effect of the iridophores, even in the wild-type, was gone after refractive index matching. This side effect of the zPACT protocol^41^may result from the combined action of the detergents present in the fixation solution and washing solution (0.1% Triton X-100, 1% DMSO, 0.1% Tween 20) and the refractive index matching solution, which itself can be slightly yellow. Together, these reagents may have washed out or diluted the pigmented compounds stored within lipid vesicles, which are responsible for the yellow colouration. This results in both phenotypes being indistinguishable from one another (Figure 15B and C).

### Development from larval to adult stage

We raised well-formed, depigmented larvae representing each phenotype to adulthood to assess their developmental viability (Figure 6) and reproductive capacity. While single-gene knock-outs exhibited an average mortality rate (from 5 dpf to 3 months of age) of <10%, *crysta-ltk* and *crystalline* fish showed higher mortality rates (∼30–40%), and *phantom* fish exhibited mortality rates >70%.

All individuals that reached adulthood were capable of normal breeding when inbred, producing offspring without abnormal mortality rates. Both *crysta-ltk/crystalline* and *slc2a11b* fish spawned F1 individuals without any obvious deleterious phenotypes or overmortality. We were also able to generate *phantom* F1 offspring by interbreeding with the same statement.

### Slc2a11 b KO does not hinder yellow pigmentation in adulthood

Unexpectedly, adult *slc2a11b* knock-out and *phantom* fish displayed yellow pigmentation, making them visually indistinguishable from wild-type fish and from *crysta-ltk* and *crystalline* individuals (Figure 13).

## Discussion

We developed multiplex gRNA sets capable of efficiently eliminating pigment formation in zebrafish larvae. The *phantom* set disrupts all three major pigment types, whereas the *crystalline* set recreates an artificial *crysta-ltk* phenotype using fewer targeted genes. However, it was difficult to distinguish the respective contributions of *mitfa* and *slc45a2* knock-outs in the *crysta-ltk* injections. The ‘slc45a2 1 and 2’ pair is likely the main contributor to both melanophore depigmentation and reduced survival, as this pair showed high efficiency and elevated malformation rates when injected alone.

Our results demonstrate that individual gRNAs can be highly efficient and may be essential for achieving complete disruption of pigment production (*tyr*, *slc2a11b*) or pigment cell differentiation (*ltk*). This is particularly important because combining multiple gRNAs in multi-gene CRISPant injections inevitably dilutes the individual gRNAs.

Increasing the overall concentration of ribonucleoprotein (RNP) complexes can partially compensate for this dilution and improve editing efficiency, however, at the expense of larval viability. We therefore selected the 1.5X concentration with two gRNAs per gene as the optimal compromise for both the *phantom* and *crystalline* gRNA sets. These sets enable PTU-free live imaging, facilitate the detection of weakly expressed fluorescent knock-ins and can reduce the time and cost of immunohistochemistry preparation by allowing direct imaging of endogenous fluorescence.

In both gRNA sets, *phantom* and *crystalline,* the major bottleneck is the *ltk* targeting strategy. The second *ltk* gRNA targets an intronic region and therefore contributes to gene disruption only when large deletions occur between the two target sites. This prompted us to investigate whether a simplified set containing only one gRNA per gene could still efficiently abolish pigmentation. However, our experiments demonstrated that targeting multiple loci per gene is generally more effective than using fewer gRNAs, although reducing the number of gRNAs improves viability. This trade-off raises concerns about off-target activity when increasing the number of gRNAs. Consequently, gRNAs should be carefully selected to minimise predicted off-target sites, particularly those without mismatches in the seed region. Although off-target cleavage is considered relatively rare when three or more mismatches are present ^42,43^, each additional gRNA introduces another potential source of unintended genomic editing. Furthermore, if stable lines are to be derived from F0 founders, predicted off-target sites should ideally not be located on the same chromosome as other targeted loci. Our results suggest that simultaneously disrupting multiple genes at several loci is challenging and may lead to large genomic rearrangements and/or increased additive off-target effects from the co-injection of multiple gRNAs (Table 2).

An alternative strategy to reduce the off-target risks associated with multiplexing would be to adapt the *phantom* gRNA set for use with LbCas12a, which has been reported to be the most efficient commercially available Cas12a variant at temperatures below 30°C ^44–47^. In addition, Cas12a can process its own crRNAs when they are separated by the appropriate direct repeat sequence ^48^. This would allow the entire gRNA set to be encoded as a single transcript, simplifying the preparation of multiplexed guide RNAs.

Regardless of the nuclease used, multiplex genome editing can induce large deletions and chromosomal rearrangements ^49–51^ and has even been associated with chromothripsis ^52^. Such genomic alterations may partly explain the moderate to high mortality rates observed in *crystalline* and *phantom* embryos, which involve four and six gRNAs, respectively.

Despite these limitations, the *phantom* and *crystalline* gRNA sets provide powerful tools for eliminating pigmentation in zebrafish larvae, facilitating live imaging, deep confocal imaging and stereomicroscopic screening of weak fluorescent signals. We are already using the *slc2a11b* knock-out to generate and identify weakly expressing green fluorescent lines that would otherwise be difficult or impossible to detect.

Interestingly, we observed that our embedding medium for confocal microscopy effectively eliminates the interference caused by yellow pigmentation and iridophores. This effect may result from refractive index matching and the slight yellow tint of the embedding medium, which could attenuate the interference associated with yellow pigmentation. However, the underlying mechanisms remain to be investigated. Nevertheless, genetic depigmentation offers the additional advantage of eliminating the need for a separate larval labelling step. This substantially reduces treatment time from 16 days to 2 days, lowers associated costs and minimises sample handling, thereby reducing the risk of physical damage.

Beyond simplifying sample preparation, this approach may also facilitate quantitative fluorescence analysis when endogenous fluorescent reporters are available. By allowing fluorescence intensity to be measured directly, it provides a readout of the fluorescent signal without the additional signal convolution associated with immunolabelling, which can arise from differences in antibody binding, labelling efficiency and staining conditions. Although immunolabelling remains valuable for detecting specific targets, these factors can complicate the quantitative assessment of target abundance. Combining genetic depigmentation with tissue clearing, therefore, offers a practical way to improve imaging efficiency while preserving sample integrity and facilitating quantitative analysis.

Based on our observations, we recommend using *crystalline* for observing red fluorescent reporters and *phantom* for detecting weak green fluorescence under a stereomicroscope, particularly when generating CRISPR-Cas9 knock-in lines. If *phantom* or *crystalline* larvae need to be raised to adulthood, we recommend using the lite versions of the respective gRNA mixes to maximise survival.

However, the benefit of the yellow-less *phantom* phenotype is lost in adulthood. Interbreeding *slc2a11b*^-/-^ mutants still produces yellow-less offspring. This recovery of yellow pigmentation in adulthood may be explained by differences between the mechanisms underlying yellow pigmentation at larval and adult stages in zebrafish and other teleosts. During larval development, pteridines are the main contributors to yellow pigmentation. Their production depends on GTP transport mediated by *slc2a11b* and subsequent processing by GTP cyclohydrolase (*gch2*) ^53,54^.

In contrast, adult yellow pigmentation mainly results from the processing and storage of dietary carotenoids within xanthophores. This pathway can be reduced by *plin6* knock-out or completely abolished by *scarb1* knock-out ^55–57^. An alternative strategy would be to target xanthophore differentiation directly, for example, through a double knock-out of *pax7a* and *pax7b*. However, this would require adding further gRNAs to the *phantom* set, potentially reducing editing efficiency and negatively affecting larval survival.

## Methods

### gRNA design

A single guide RNA (gRNA) targeting the third exon of the *alpk1* gene was designed using CRISPOR (http://crispor.tefor.net/) against the *Danio rerio* UCSC May 2017 (GRCz11/danRer11) reference genome. The gRNA was optimised for SpCas9, SpCas9-HF1 and eSpCas9 1.1 enzymes using the 20bp-NGG PAM sequence. Selection criteria included proximity to the target locus, high specificity score, minimal predicted off-target effects and absence of sequence motifs that could compromise cleavage efficiency.

The gRNA was assembled by annealing 100 μM crRNA (sequence: gRNA 1, gRNA 2) with 100 μM tracrRNA (Integrated DNA Technologies, IDT). The annealing protocol consisted of heating at 95°C for 5 minutes, followed by gradual cooling to room temperature (20°C). The annealed gRNA was diluted in IDT duplex buffer to a working concentration of 20 μM.

Primers were designed on Primer-BLAST, National Library of Medicine (online: https://www.ncbi.nlm.nih.gov/tools/primer-blast/index.cgi).

### Microinjections

CRISPR-Cas9 gene editing was performed using SpCas9 HiFi (Museum National d’Histoire Naturelle) ribonucleoprotein (RNP) complexes. The concentration of gRNA and Cas9 was optimised for each injection condition: 13.33 µM gRNA and 10 µM Cas9 for single gRNA injections, 6.67 µM gRNA and 5 µM Cas9 for duplex injections, 2.22 µM gRNA and 1.67 µM Cas9 for double CRISPants *phantom* injections and 4.44 µM gRNA and 3.34 µM Cas9 for single CRISPants *phantom* injections. crRNA and tracrRNA (Integrated DNA Technologies) were denatured and annealed by heating at 95°C for 5 minutes, followed by slow cooling to room temperature (25°C) to form the gRNA duplex. Cas9 and gRNAs were then co-incubated at 37°C for 10 minutes to allow RNP complex formation. For microinjection, RNP complexes were loaded into heat-pulled glass capillaries (Harvard Apparatus, dimensions: 1.0 OD x 0.58 ID x 100 L) and injected into one-cell stage embryos. Embryos were aligned in a 1.5% agarose-embryo medium mould, stabilising the eggs during the injection procedure, with the embryo’s lateral surface exposed for microinjection. After injection, embryos were placed in a 28°C incubator for development and phenotypic analysis.

### DNA extraction and sequence analysis

Genomic extractions were performed with the Rapid Lyse extraction kit from Macherey-Nagel at 24 hours post fertilisation (hpf), prior to the first appearance of pigmentation.

PCR was performed with GoTaq G2 DNA Polymerase Hot Start from Promega (5X GoTaq Reaction Buffer, MgCL2 1,5 mM, dNTPs 10 mM, forward and reverse primers 10 µM each, GoTaq G2 DNA Polymerase Hot Start 1,25u, template DNA <0,5µg/µL).

A gel of 1,5% agarose in 0,5X TBE was made to verify PCR amplifications with GelRed at 2,5% to reveal the bands.

Validated amplicons were sent to Eurofins Genomics for Sanger sequencing to retrieve the chromatograms, then analysed by ICE online software developed by Synthego (online: https://ice.editco.bio/#/).

### Raising conditions for larvae

Injected and non-injected zebrafish embryos (Danio rerio) were maintained in E3 medium containing 5 mM NaCl, 0.17 mM KCl, 0.33 mM CaCl₂ and 0.33 mM MgSO₄ in deionized water. The pH was adjusted to 7.2. Methylene blue was added for 24 hours at 28,5°C to be then washed with clear E3 medium and incubated at 28,5°C until 5 dpf. Each petri dish contained 50 embryos regardless of the conditions of injection.

### Zebrafish Housing and Husbandry

Adult zebrafish were housed at a density of five individuals per litre in controlled aquatic conditions, in compliance with European standards for zebrafish maintenance and welfare. The following environmental parameters were maintained: a 14:10 h light-dark photoperiod, water temperature at 26.5 ± 1 °C, pH maintained at 7.8 ± 0.1, conductivity at 240 ± 30 μS/cm, with NH₄⁺ and NO₂⁻ levels below detection limits and NO₃⁻ concentrations kept under 50 mg/L.

Fertilised embryos were maintained at 28 °C until 5 dpf. For the initial two-week feeding period, larvae received live rotifers (*Brachionus plicatilis*) at a rate exceeding 500 individuals per fish daily, after which they were transitioned to commercial dry feed (Skretting, Gemma Micro) administered twice daily until apparent satiation.

The experimental work utilised the AB zebrafish strain (ZFIN_ZDB-GENO-960809-7) sourced from the European Zebrafish Resource Centre, Germany (ZDB-LAB-130607-1). All experimental protocols adhered to European Union Directive 2011/63/EU requirements and received approval from the institutional ethics committee (authorisation no. 59 CEEA).

### Zebrafish KO-lines generated in this study

This study generated the following new knock-out lines: *crysta-ltk*: Tg(); *crystalline*: Tg(); *crystalline-lite*: Tg(); *phantom*: Tg(); *phantom-lite*: Tg().

### Animals and fixation

Zebrafish larvae at 5 dpf were anaesthetised with tricaine (MS-222; 150 mg/L buffered with sodium bicarbonate) and subsequently euthanised by gradual exposure to an overdose of tricaine. Larvae were then fixed for 24 h at 4°C in freshly prepared fixative containing 1× phosphate-buffered saline (PBS), 4% paraformaldehyde (PFA), 0.1% Triton X-100 and 1% dimethyl sulfoxide (DMSO). Following fixation, samples were washed for 24 h at room temperature (RT) in PBST (1× PBS containing 0.1% Tween 20) supplemented with 0.05% sodium azide.

### Depigmentation

Fixed larvae were washed in PBST for 1-3 h at RT and subsequently incubated for 2 h at RT in freshly prepared depigmentation solution containing 1× PBS, 5% hydrogen peroxide (H₂O₂), 0.1% Triton X-100 and 0.05% sodium azide. Depigmentation was performed under intense illumination. Samples were then rinsed three times with PBST and washed overnight in PBST at RT.

### Fluorescent staining

Samples were incubated for 6 h at RT with gentle agitation in blocking solution containing 10% normal goat serum (NGS), 10% DMSO, 100 mM PBS–glycine, 0.5% Triton X-100, 1× PBS and 0.1% Tween 20. Cell membranes and double-stranded DNA were fluorescently labelled using the lipophilic tracer DiI C18(3) (Invitrogen, Thermo Fisher Scientific; Cat. No. D282) and the fluorescent nucleic acid stain YOYO-1 iodide (Thermo Fisher Scientific; Cat. No. Y3601), respectively. Samples were incubated for 48 h at RT in a staining solution containing YOYO-1 iodide and DiI at a final concentration of 0,05 µg/µL and 1 µg/µL, respectively. The staining solution consisted of 2% NGS, 10% DMSO, 0.05% sodium azide, 0.2% Triton X-100, 1× PBS, 0.1% Tween 20 and 200 mM NaCl. Samples were protected from light during staining. Following incubation, samples were washed at least three times for 20 min each in PBST.

### Refractive index matching

To match the refractive index of the samples to that of the imaging medium, larvae were incubated in a 1:1 mixture of MD+ clearing solution (MD+)^58^ and PBST for at least 5 h at RT, followed by incubation in 100% MD+ for at least 5 h at RT. MD+ clearing solution consisted of 45% (w/v) sucrose, 20% (w/v) nicotinamide, 10% (w/v) triethanolamine and 0.1% Triton X-100.

### Mounting

A 1.5% (w/v) agarose solution prepared in a mixture of water and MD+ clearing solution was poured into a Petri dish. The final mixture contained 30% water and 70% MD+, with the composition adjusted according to the sample requirements. A custom-designed, 3D-printed mould was used to create wells in the agarose gel. Samples were positioned in the wells with their ventral sides facing downwards, and the wells were sealed with a thin layer of agarose solution. The mounted samples were immersed in MD+ and incubated for 24 hours at RT in the dark to facilitate refractive index equilibration by diffusion.

### Confocal imaging

Image acquisition was performed using a Nikon A1R HD resonant-scanning laser confocal microscope equipped with a CFI Plan Apo 10×C Glyc objective with a 5.5-mm working distance, a piezoelectric positioning system, 488-nm and 561-nm diode lasers and gallium arsenide phosphide (GaAsP) detectors. Images were acquired at a resolution of 1024 × 1024 pixels, with a Z-step size of 0.8 µm. Mineral oil was applied to the surface of the imaging medium to minimise evaporation and maintain a stable refractive index during image acquisition.

### Software

Editing and annotations of DNA and protein sequences were performed with SerialCloner^59^. DNA sequencing chromatograms were inspected and analysed using FinchTV ^60^. Confocal microscopy images were acquired using NIS-Elements ^61^. Microscopy images were processed and analysed using Fiji ^62^. Image preparation and figure assembly were performed using GIMP ^63^ and Inkscape ^64^.

During manuscript preparation, the authors used artificial intelligence (AI) to assist with language editing, sentence reformulation and improvements in clarity and readability. Grammarly ^65^ was used for spelling, grammar and stylistic corrections. All AI-assisted suggestions were critically reviewed and, where necessary, revised by the authors, who take full responsibility for the final manuscript content.

## Supporting information

All supplementary material

## ACKNOWLEDGMENTS

We are sincerely grateful to Jean-Paul Concordet for kindly providing the recombinant Cas9 protein used in this study, as well as for his valuable technical advice and insightful discussions. We also gratefully acknowledge the support of the TACGENE platform at the Museum National d’Histoire Naturelle (MNHN; CNRS UMR 7196, INSERM U1154). The authors also thank Marie-Elise Schwartz, facility manager, Théo Perrot, Noé Sauger, Gaël Mollat and Ronan Mudet for their technical expertise and assistance with fish husbandry, animal care and welfare monitoring.

We thank Muriel Perron and Albert Chesneau for their contribution in finding the strategy for generating the CRISPants.

## Contributions

A.B, P.A. and J-P.L. designed research; A.B., P.A. and M.J. performed research; A.B., P.A., M.J. and J-P.L analysed data; A.B. and A.J. prepared all figures; A.B., A.J., P.A., M.J. and J-P.L wrote the manuscript.

## DATA AVAILABILITY

The original data supporting the findings of this study have been deposited in the Zenodo repository and are publicly available at https://doi.org/10.5281/zenodo.23660844.

