## Supplementary material for "I. Phantom: a multiplex CRISPR-Cas9 gRNA toolkit for efficient generation of fully unpigmented F0 zebrafish": All supplementary material

### Supplemental Material

| Gene | gRNA | Sequence | Cut position | Gene length |
|---|---|---|---|---|
| *ltk* | 1 | CAGAACATTGTCCGCTGCAT | Exon 22 | 28 exons |
| *ENSDART00000061547.7* | 2 | TGGCCTACAGTACATGGTCC | Intron 22 | |
| | 3 | TAGGACCCAGGAACCGTAGG | Exon 28 | |
| *mitfa* | 1 | CTTCGAGGGGGTTTCCAGGT | Exon 2 | 9 exons |
| *ENSDART00000056457.5* | 2 | GGGGTCATGCCGTGCTCGGC | Exon 2 | |
| | 3 | CACGGCATGACCCCGGGACC | Exon 2 | |
| | 4 | TTGAGGGTGAGAAGGGCCAT | Exon 2 | |
| | 5 | ATGTGCATCATGCCCGGAGA | Exon 3 | |
| | 6 | ATGGACAAAGCTGGACCATG | Exon 3 | |
| *slc45a2* | 1 | GACCGTACATACTCTTACTG | Exon 1 | 7 exons |
| *ENSDART00000030887.8* | 2 | CTTTAAGCTCCGAGTCCACC | Exon 1 | |
| | 3 | AAGGACATGGGCCATCGTGG | Exon 2 | |
| *tyr* | 1 | ACCTGACTGGACGCCGCACA | Exon 1 | 5 exons |
| *ENSDART00000122238.3* | 2 | TGGTGTCCGACCTTCCCAAC | Exon 1 | |
| *slc2a11b* | 1 | GTCATTGGACGCTTCCTGAC | Exon 4 | 12 exons |
| *ENSDART00000092333.6* | 2 | GAGGAGGGTTAACATATCAC | Exon 3 | |

***Table 1: Tested gRNAs and their localisation in their corresponding gene.***

*For each gene, a transcript was chosen as the main transcript to knock-out. The sequence for each gRNA is detailed, as well as its location in the gene relative to its length.*

| Gene | Chr | sgRNA | Sequence | Off-targets mismatches | | | | | | | | | | | |
|---|---|---|---|---|---|---|---|---|---|---|---|---|---|---|---|
| | | | | Whole sequence | | | | | | None in seed | | | | | |
| | | | | 0 | 1 | 2 | 3 | 4 | Sum | 0 | 1 | 2 | 3 | 4 | Sum |
| *ltk* | 17 | 1 | CAGAACATTGTCCGCTGCAT | 0 | 0 | 0 | 3 | 16 | 19 | 0 | 0 | 0 | 0 | 0 | 0 |
| | | 2 | TGGCCTACAGTACATGGTCC | 0 | 0 | 0 | 3 | 27 | 30 | 0 | 0 | 0 | 0 | 1 | 1 |
| | | 3 | TAGGACCCAGGAACCGTAGG | 0 | 0 | 0 | 0 | 5 | 5 | 0 | 0 | 0 | 0 | 0 | 0 |
| *mitfa* | 6 | 1 | CTTCGAGGGGGTTTCCAGGT | 0 | 0 | 0 | 3 | 10 | 13 | 0 | 0 | 0 | 1 | 0 | 1 |
| | | 2 | GGGGTCATGCCGTGCTCGGC | 0 | 0 | 0 | 1 | 6 | 7 | 0 | 0 | 0 | 0 | 0 | 0 |
| | | 3 | CACGGCATGACCCCGGGACC | 0 | 0 | 0 | 0 | 3 | 3 | 0 | 0 | 0 | 0 | 0 | 0 |
| | | 4 | TTGAGGGTGAGAAGGGCCAT | 0 | 0 | 0 | 2 | 13 | 15 | 0 | 0 | 0 | 0 | 0 | 0 |
| | | 5 | ATGTGCATCATGCCCGGAGA | 0 | 0 | 0 | 2 | 41 | 43 | 0 | 0 | 0 | 0 | 1 | 1 |
| | | 6 | ATGGACAAAGCTGGACCATG | 0 | 0 | 0 | 1 | 55 | 56 | 0 | 0 | 0 | 0 | 0 | 0 |
| slc45a2 | 21 | 1 | GACCGTACATACTCTTACTG | 0 | 0 | 0 | 1 | 8 | 9 | 0 | 0 | 0 | 0 | 0 | 0 |
| | | 2 | AAGGACATGGGCCATCGTGG | 0 | 0 | 0 | 0 | 9 | 9 | 0 | 0 | 0 | 0 | 0 | 0 |
| | | 3 | CTTTAAGCTCCGAGTCCACC | 0 | 0 | 0 | 0 | 6 | 6 | 0 | 0 | 0 | 0 | 0 | 0 |
| *tyr* | 15 | 1 | ACCTGACTGGACGCCGCACA | 0 | 0 | 1 | 1 | 4 | 6 | 0 | 0 | 0 | 0 | 0 | 0 |
| | | 2 | TGGTGTCCGACCTTCCCAAC | 0 | 0 | 0 | 2 | 9 | 11 | 0 | 0 | 0 | 0 | 0 | 0 |
| *slc2a11b* | 5 | 1 | GTCATTGGACGCTTCCTGAC | 0 | 0 | 0 | 0 | 18 | 18 | 0 | 0 | 0 | 0 | 0 | 0 |
| | | 2 | GAGGAGGGTTAACATATCAC | 0 | 0 | 0 | 1 | 25 | 26 | 0 | 0 | 0 | 0 | 2 | 2 |

***Table 2: Tested gRNAs and their respective off-targets with and without mismatches in the seed region of their sequence.***

*Off-target sites were divided into two categories: sequences with 0–4 mismatches relative to the target, and a subset where mismatches occur exclusively in the PAM-distal, non-seed region (positions 13–20), leaving the seed region (positions 1–12) intact. These sites are expected to carry the highest risk of unwanted cleavage, with risk proportional to the number of off-targets. The risk is proportional to the number of off-targets.*

| Line \ Gene | *ltk* | | *mitfa* | | | | | *slc45a2* | | *tyr* | *slc2a11b* |
|---|---|---|---|---|---|---|---|---|---|---|---|
| gRNA ID | 2 | 3 | 2 | 3 | 4 | 5 | 6 | 2 | 3 | 2 | 2 |
| *ltk 1* | + | + | | | | | | | | | |
| *ltk 2* | - | + | | | | | | | | | |
| *mitfa 1* | | | + | + | + | - | - | | | | |
| *mitfa 2* | | | - | + | + | - | - | | | | |
| *mitfa 3* | | | - | - | + | - | - | | | | |
| *mitfa 4* | | | - | - | - | + | + | | | | |
| *slc45a2 1* | | | | | | | | + | + | | |
| *slc45a2 2* | | | | | | | | - | + | | |
| *tyr 1* | | | | | | | | | | + | |
| *slc2a1b 1* | | | | | | | | | | | + |

***Table 3: Combinations of gRNA constructs***

*To enhance depigmentation by knocking-out each gene, we designed pairs of gRNAs. The tested gRNA combinations are indicated by a "+".*

| Line \ Gene | *ltk* | | *mitfa* | | *slc45a2* | | *tyr* | | *slc2a11b* | |
|---|---|---|---|---|---|---|---|---|---|---|
| gRNA ID | 1 | 2 | 4 | 5 | 1 | 2 | 1 | 2 | 1 | 2 |
| *crysta-ltk* | + | + | + | + | + | + | - | - | - | - |
| *crystalline* | + | + | - | - | - | - | + | + | - | - |
| *crystalline-lite* | + | - | - | - | - | - | + | - | - | - |
| *phantom* | + | + | - | - | - | - | + | + | + | + |
| *phantom-lite* | + | - | - | - | - | - | + | - | + | - |

***Table 4: Composition of each mix of gRNA.***

*Genes targeted by each gRNA mix, and their corresponding gRNA IDs chosen for each gene, are indicated with a "+".*

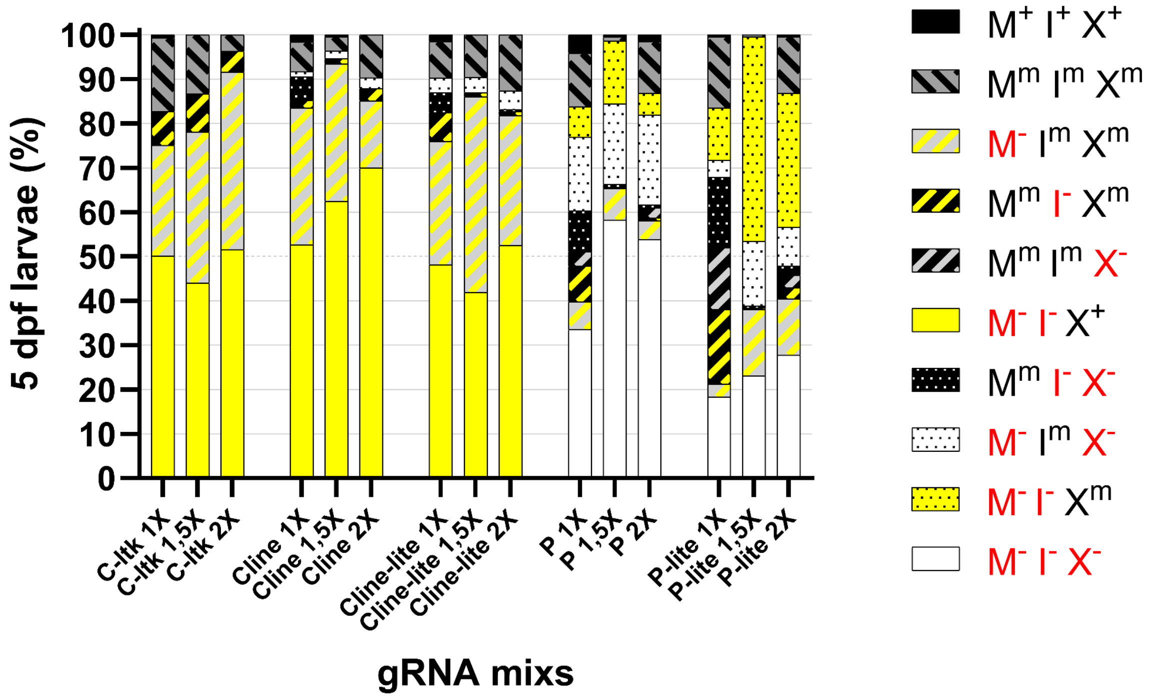


***Figure 1: (supplementary 1). Detailed proportions of each phenotype for each pigment at 5 days post-depigmentation injections.***

*C-ltk (crysta-ltk), Cline (crystalline), Cline-lite (crystalline-lite), P (phantom) and P-lite (phantom-lite) gRNA sets were injected into three batches of 100 single-cell embryos at 1X, 1.5X and 2X concentrations. Every phenotype is reported in MIX categories (M: melanophores, I: iridophores, X: xanthophores; +: fully pigmented, m: mosaic, -: unpigmented), and the number of individuals per category is displayed.*

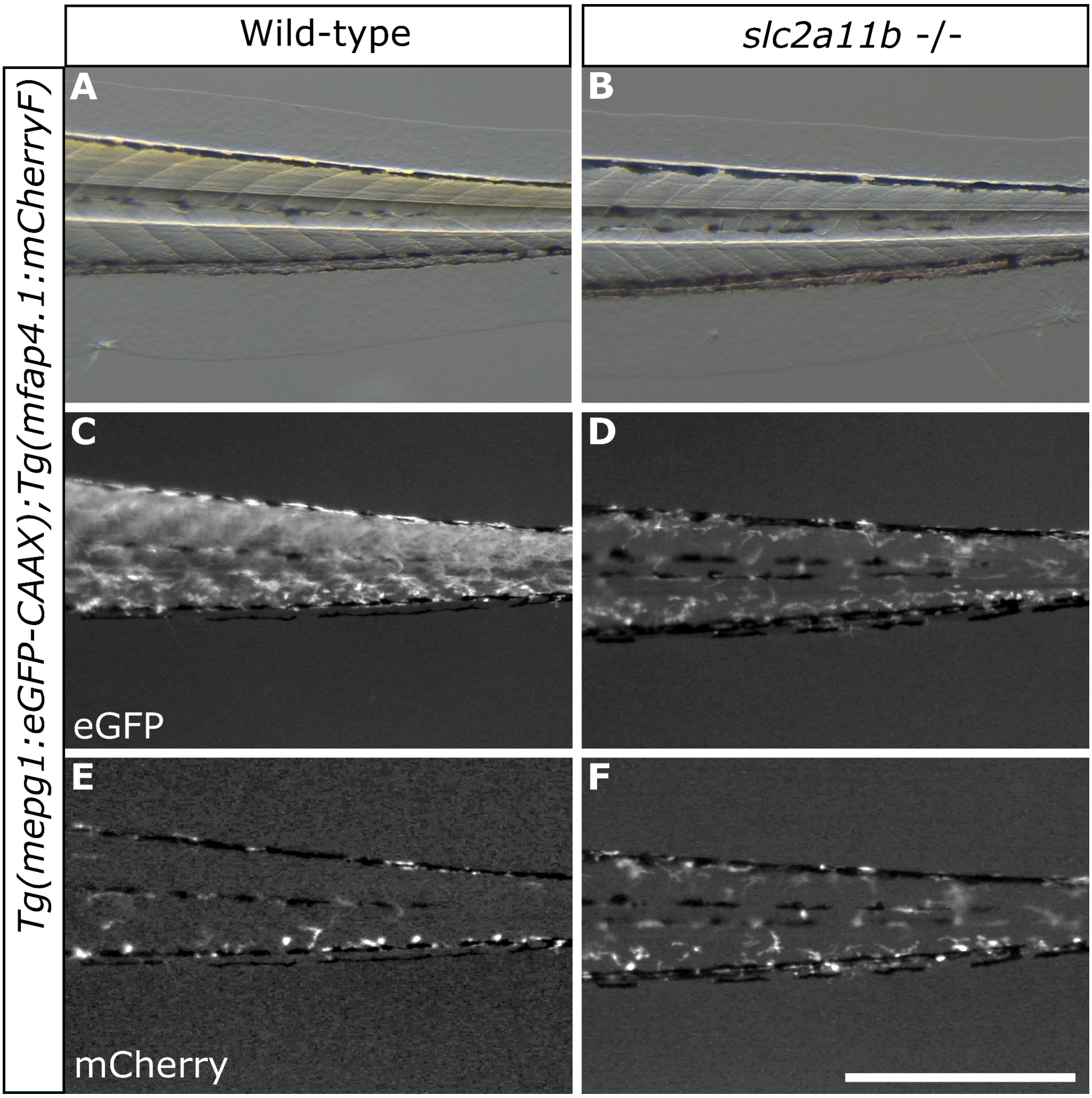


***Figure 2: (supplementary 2). Qualitative comparison between Tg(mpeg1:eGFP-CAAX);Tg(mfap4.1:mCherryF) zebrafish injected with slc2a11b-targeting gRNAs or non-injected (tails).***

*The Tg(mpeg1:eGFP-CAAX) line was crossed with the Tg(mfap4.1:mCherryF) line to generate double transgenic embryos, which were either injected at the 1-cell stage with slc2a11b* ***(B, D, F)*** *or left non-injected as wild-type controls* ***(A, C, E)****. At 5 dpf, larvae displaying expected wild-type and slc2a11b phenotypes were sorted and imaged under a fluorescence stereo microscope to assess xanthophore autofluorescence interference with macrophage pattern visibility in both green* ***(eGFP-CAAX, C and D)*** *and red* ***(mCherry, E and F)*** *channels. Brightness and contrast were uniformly optimised for both green and red channel images. Scale bar: 0.5 mm.*

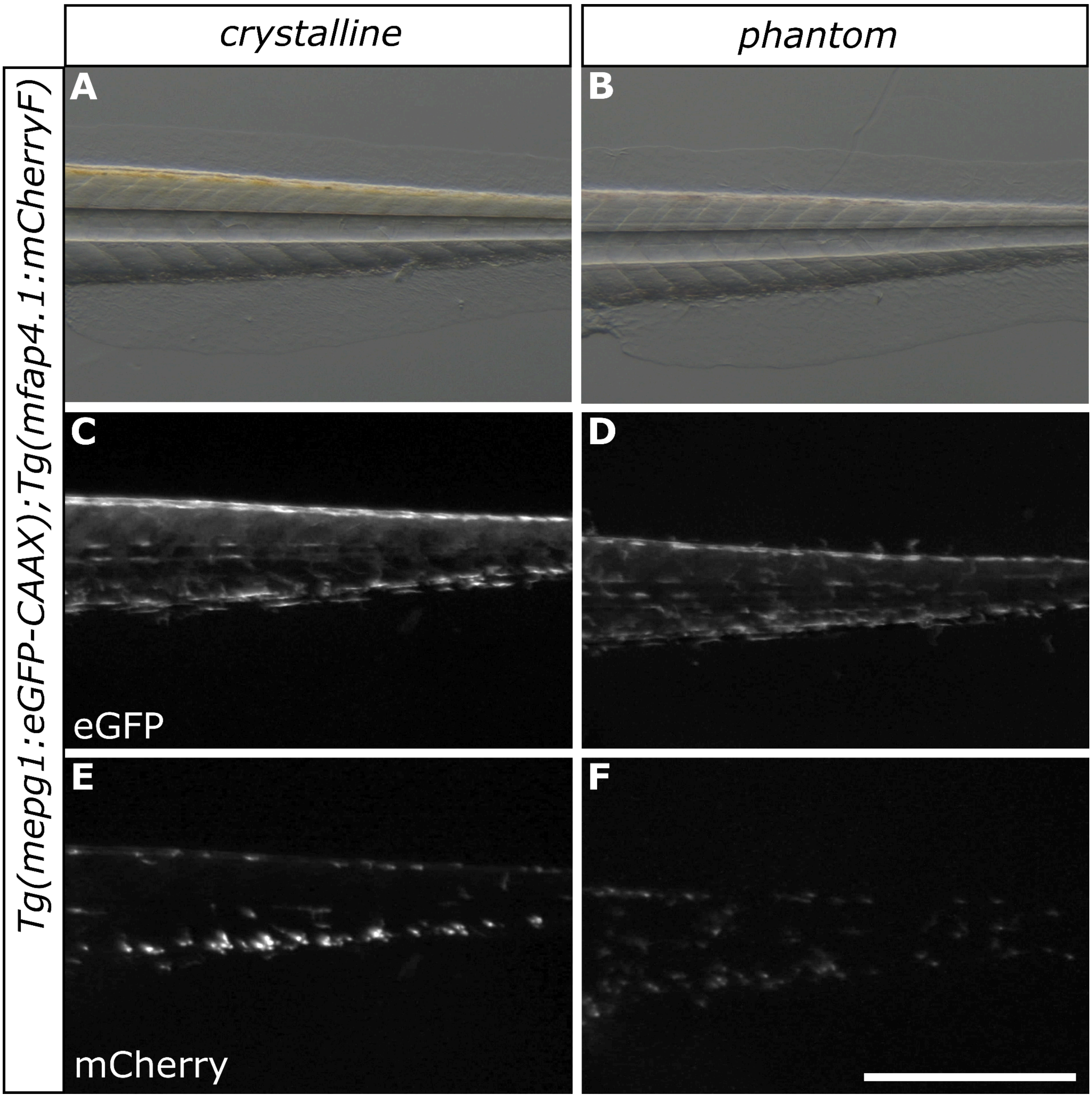


***Figure 3: (supplementary 3): qualitative comparison between Tg(mpeg1:eGFP-CAAX);Tg(mfap4.1:mCherryF) injected with crystalline or phantom gRNA sets (tails).***

*The Tg(mpeg1:eGFP-CAAX) line was crossed with the Tg(mfap4.1:mCherryF) line to generate double transgenic embryos injected at the 1-cell stage with either crystalline (A, C, E) or the phantom gRNA set (B, D, F). Larvae showing the expected crystalline or phantom phenotypes were sorted and imaged under a fluorescence stereo microscope at 5 dpf to show the effect of xanthophore autofluorescence on the visibility of the macrophage pattern, both in green (Green) (C and D) and red (mCherry) (E and F). Brightness and contrast were uniformly optimised for both green and red channel images. Scale bar: 0.5 mm.*

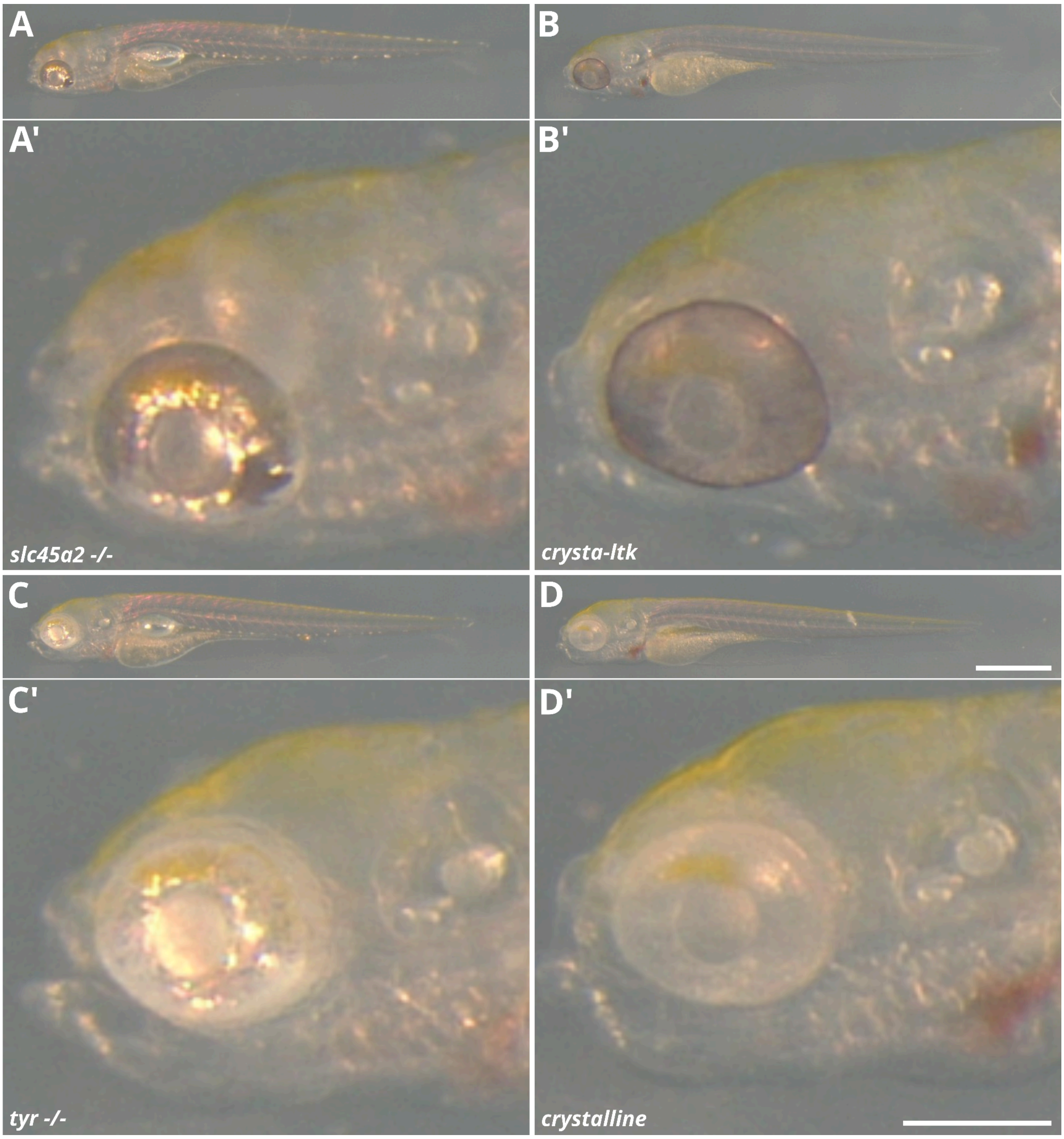


***Figure 4: (supplementary 4). Magnification on the heads of slc45a2 and tyr albino phenotypes and their crystal-related phenotype:***

***(A)*** *slc45a2: The eye remains slightly pigmented beneath the reflective iridophore layer.* ***(B)*** *crysta-ltk: The blackness of the eye without the iridophore layer is more visible than in the slc45a2 knock-out.* ***(C)*** *tyr: Only the iridophore layer is visible in the eye.* ***(D)*** *crystalline: The back of the right eye is visible behind the totally unpigmented left eye, with yellow xanthophores clearly observable in or behind the left eye. Brightness and contrast were uniformly optimised for all images. Scale bar: 0.5 mm.*

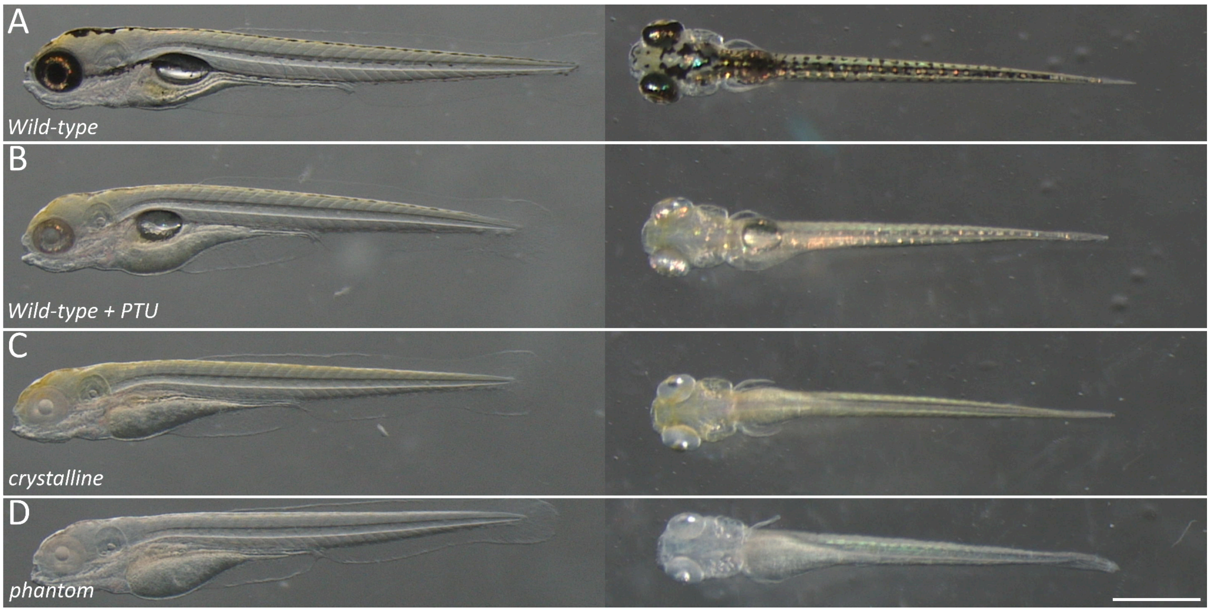


***Figure 5: (supplementary 5). Comparison of transparency between wild-type, PTU-treated wild-type, crystalline and phantom phenotypes in lateral and dorsal views.***

*5 dpf larvae were sorted for expected phenotypes under angled illumination from above to visualise iridophore shine (orange spots) under a stereo microscope. Wild-types were non-injected* ***(A)****, while a subset of the same wild-type population was incubated with 0.2 mM PTU at 1 dpf to inhibit melanophore pigmentation* ***(B)****. Additional wild-types from the same spawn were injected with crystalline* ***(C)*** *or phantom* ***(D)*** *gRNA sets at 1.5X. Scale bar: 0.5 mm*

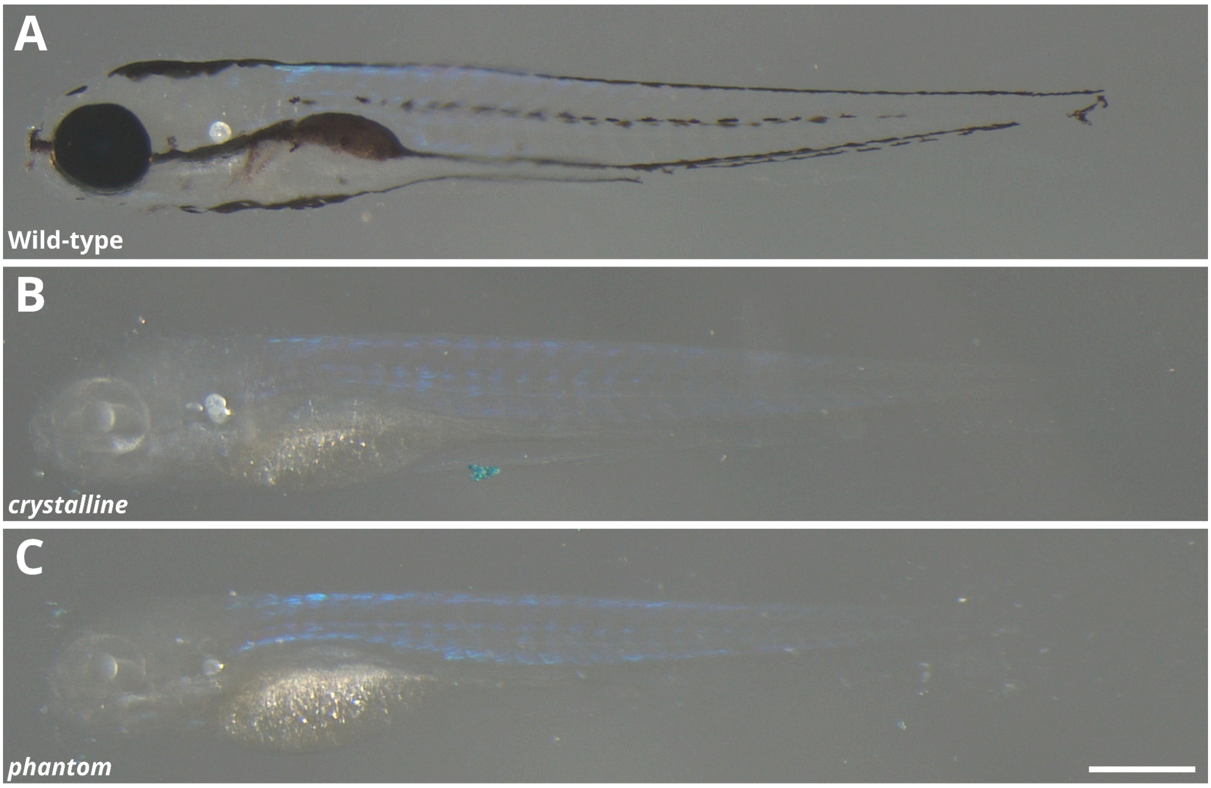


***Figure 6: (supplementary 6). Effects of tissue clearing on endogenous pigmentation.***
*5 dpf larvae were treated according to the zPACT protocol to enable imaging. Yellow pigmentation is no longer visible in any larva, nor are iridophores in wild-type* ***(A)****, due to refractive index matching. The yellow pigmentation is likely lost due to a combination of weak detergent (Tween 20) and the slightly yellow clearing solution, which may have washed away or diluted the components responsible for this pigmentation. Consequently, crystalline* ***(B)*** *and phantom* ***(C)*** *phenotypes become indistinguishable from one another. Scale bar: 0.5 mm.*

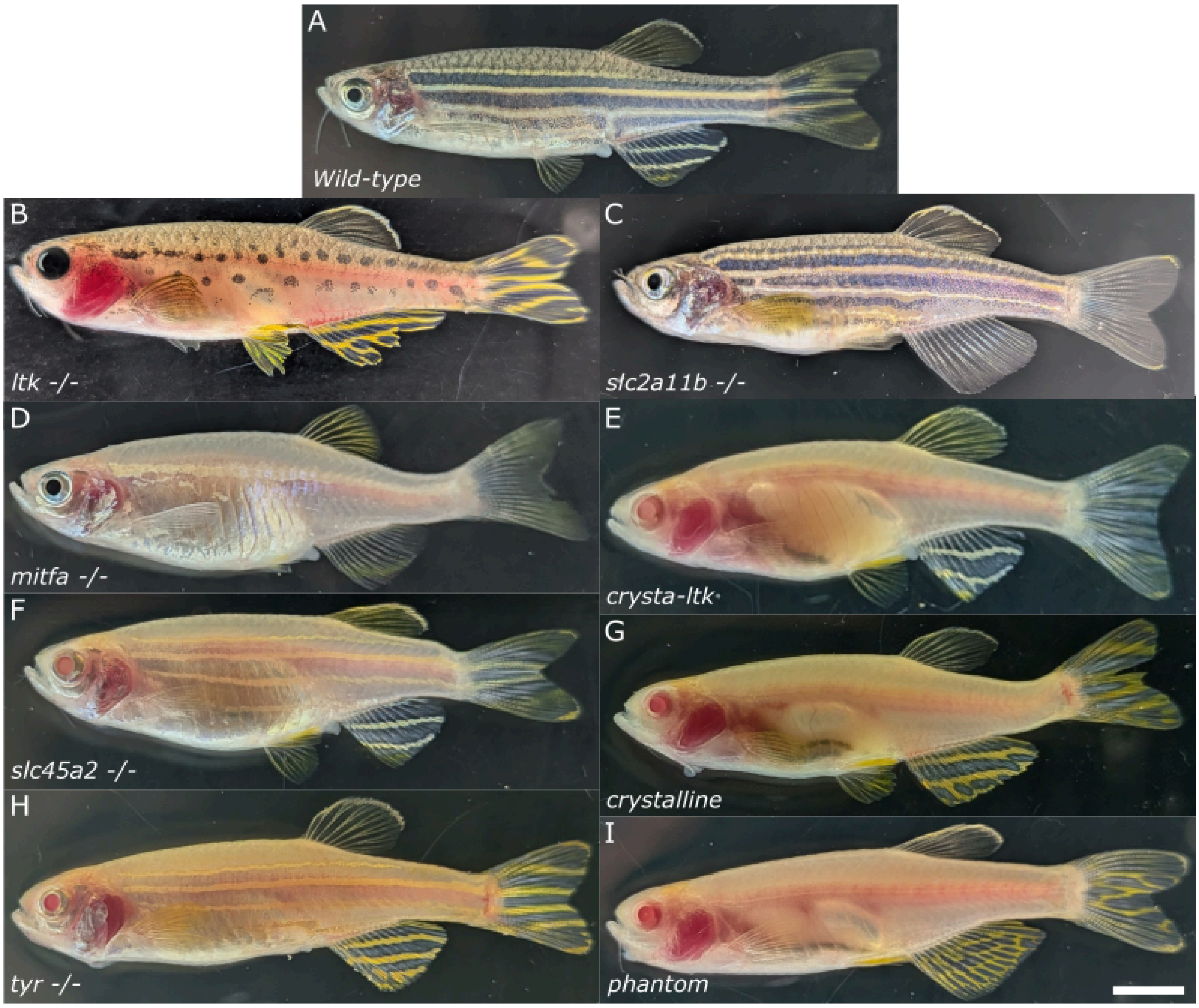


***Figure 7: (supplementary 7). Illustration of pigmentless phenotypes in F0 adult fish.***

*All 5 dpf larval phenotypes described in this study were raised to adulthood without visible defects or breeding incapacity. Eye pigmentation differences observed at larval stages between slc45a2⁻/⁻ and tyr⁻/⁻ are no longer visible. slc2a11b⁻/⁻ and phantom adults display wild-type-like yellow pigmentation despite its absence at the larval stage. Scale bar: 0.5 cm.*
